# Single-cell genomics of single soil aggregates: methodological assessment and potential implications with a focus on nitrogen metabolism

**DOI:** 10.1101/2025.01.09.632267

**Authors:** Emi Matsumura, Hiromi Kato, Shintaro Hara, Tsubasa Ohbayashi, Koji Ito, Ryo Shingubara, Tomoya Kawakami, Satoshi Mitsunobu, Tatsuya Saeki, Soichiro Tsuda, Kiwamu Minamisawa, Rota Wagai

**Affiliations:** Institute for Agro-Environmental Sciences (NIAES), National Agriculture and Food Research Organization (NARO), Ibaraki, Japan; Graduate School of Life Science, Tohoku University, Miyagi, Japan; Research Center for Advanced Analysis (NAAC), National Agriculture and Food Research Organization (NARO), Ibaraki, Japan; Graduate School of Agriculture, Ehime University, Ehime, Japan; bitBiome Inc., Tokyo, Japan

**Keywords:** water-stable macroaggregates, soil microbial ecology, cell extraction method, sonication, nitrogen cycling, bacterial community compositions, functional diversity, EPS related gene

## Abstract

Soil particles in plant rooting zone are largely clustered to form complex porous structural unit called aggregates where highly diverse microbes coexist and drive biogeochemical cycling. The complete extraction of microbial cells and DNA from soil is a substantial task as certain microbes exhibit strong adhesion to soil surfaces and/or inhabit deep within aggregates. Yet, the degree of aggregate dispersion and the efficacy of extraction have rarely been examined, and thus adequate cell extraction method from soil remain unclear. We aimed to develop an optimal method of cell extraction for single-cell genomics (SCG) analysis of single soil aggregates by focusing on water-stable macroaggregates (diameter: 5.6-8.2 mm) isolated from a topsoil of cultivated Acrisol. Using the same six individual aggregates, we performed both SCG sequencing and amplicon analysis. While both bead-vortexing and sonication dispersion methods improved the extractability of bacterial cells compared to previous studies, the latter yielded higher number and more diverse microbes compared to the former. The analyses of nitrogen-cycling and exopolysaccharides-related genes suggested that the sonication-assisted extraction led to greater recovery of microbes strongly attached to soil particles and/or inhabited the aggregate subunits that were more physically stable (e.g., aggregate core). Further SCG analysis revealed that all six aggregates held intact microbes having the genes (i.e., potentials) to convert nitrate into all possible nitrogen forms while some low-abundance genes showed inter-aggregate heterogeneity. In addition, all six aggregates studied showed overall similarity in pore characteristics, phylum-level composition, and the microbial functional redundancy. Together, these results suggest that water-stable macroaggregates may act as a functional unit in soil and show potential as a useful experimental unit in soil microbial ecology. Our study also suggest that conventional methods employed for the extraction of cell and DNA may not be optimal. The current findings underscore the necessity to advance extraction methodologies, thereby facilitating a more comprehensive understanding of the microbial diversity and functioning within soil environments.

## 1 Introduction

Soil microorganisms inhabit a highly heterogeneous environment that varies both spatially and temporally, driving biogeochemical cycles essential to terrestrial ecosystems. Although microbial communities are highly diverse (Gans et al., 2005), most biogeochemical transformations are mediated by a limited set of metabolic pathways shared across various taxonomic groups (Louca et al., 2018). Among these cycles, the nitrogen (N)-cycling is tightly controlled by specific groups of soil microbes (Hayatsu et al., 2008, Levy-Booth et al., 2014, Kuypers et al., 2018). Nitrous oxide (N₂O) is a potent greenhouse gas and ozone-depleting substance (Ravishankara et al., 2009). The production and transformation of N₂O in the soil is controlled by microbial communities, soil properties, and environmental conditions (Wan et al., 2024, Wrage-Mönnig et al., 2018, Zhuang et al., 2024). Denitrification and N₂O reduction basically occur under anaerobic conditions (Schlüter et al., 2024). In the final step of the denitrification process (N₂O → N₂), the primary taxa of *nosZ*-harboring bacteria responsible for nitrous oxide reduction in soil include Alpha-proteobacteria (e.g., Rhizobiaceae), Beta-proteobacteria (e.g., Burkholderiales), Bacteroidota, Gemmatimonadota, and Delta-proteobacteria (Hallin et al., 2018). However, much remains to be understood. Gaining insights into the ecology of nitrogen-cycle-associated microorganisms in soil is crucial for a comprehensive understanding of N₂O production mechanisms in agricultural soils.

Soil structure, in particular aggregates, play a critical role in the creation of heterogeneous microenvironments that are physically stable habitat for microorganisms compared to their surrounding soil matrix (Oades and Waters, 1991). Some aggregates are distinct in surface soils and can be separated by water-sieving as water-stable aggregates (Oades and Waters, 1991). A porous soil-aggregate are formed as a consequence of development of the surface soil layer in pedogenes (Totsche et al., 2018). In a conceptual model of aggregate hierarchy, it has been proposed that the small particles including microaggregate (< 250 µm) are bound together by ephemeral binding agents (e.g. fine roots and fungal hyphae) in macro-aggregate, defined as > 250 µm diameter (Tisdall and Oades, 1982). Conversely, the micro-aggregate exhibits greater physical stability due to the strong binding of iron (hydr) oxide and short-range order minerals via microbial debris and extracellular polymeric substances (Chenu and Cosentino, 2011, Six et al., 2004, Totsche et al., 2018). The diverse physicochemical properties of the soil solid phase within the hierarchic structure contribute to form the heterogeneous microenvironments (Gupta and Germida, 2015), and they can be isolated by dispersion at progressively increasing energy level (Kaiser and Asefaw Berhe, 2014). Good soil structure, which relies on effective aggregation, is an integral part of ecosystem functioning (Rabot et al., 2018, Costa et al., 2018, Bronick and Lal, 2005).

The heterogeneity of soil matrix contributes to the distribution of distinct microbial community compositions, which may vary according to aggregate size (Sessitsch et al., 2001, Hemkemeyer et al., 2015) or the interior and exterior of aggregates (Ranjard et al., 1998, Hattori, 1967). Recent research by (Mitsunobu et al., 2025) has shown that the dynamics of microbial N₂O functional groups and O₂/N₂O concentrations are regulated by soil microstructure (e.g., pore networks) using a detailed anatomical approach to single soil aggregates. This study suggests that the physical structure at the scale of individual aggregates influences the habitats of N₂O-reducing bacteria. Specifically, it was observed that *nosZ*-harboring bacteria were localized deep within the interiors of macroaggregates. However, fundamental questions remain regarding the interactions between soil structural heterogeneity and microbial function. For instance, although aggregates can be considered relatively discrete physical units that maintain natural soil structure, the degree to which microbial community composition varies across individual aggregates is still unknown. Key questions include: What determines the microbial community structure within individual aggregates? What determines the microbial community structure within individual aggregates? What are the taxonomic identities and genetic profiles of bacteria within these aggregates, and where are they specifically located within the aggregate matrix? Addressing these questions will require innovative methods beyond conventional bulk soil genome analysis, which typically assumes soil is a homogeneous environment.

Advances in molecular biology have made metagenomics a powerful tool for studying the relationship between functional and taxonomic diversity in microbial communities (Tringe et al., 2005, Pan et al., 2014, Leff et al., 2015, Fierer et al., 2012b, Fierer et al., 2012a, Fierer et al., 2013). The two primary metagenomic methods are amplicon analysis and shotgun sequencing. Shotgun metagenomics, which provides both functional and taxonomic insights, has several advantages and limitations. One major advantage of shotgun metagenomics is that it allows the abundance of each gene to be associated with specific ecological processes, enabling the simultaneous examination of multiple ecosystem functions within a single soil sample (Allison and Martiny, 2008). This multifaceted approach recognizes the significance of ecosystem multifunctionality (Hector and Bagchi, 2007). However, shotgun metagenomics also has some limitations. For example, ribosomal protein genes are often absent from metagenome-assembled genomes (MAGs), complicating functional profiling (Mise and Iwasaki, 2022). Moreover, metagenomic sequencing struggles to achieve strain-resolved genomes (Arikawa and Hosokawa, 2023), despite the substantial functional diversity that can exist between strains (Lin et al., 2016, Krause et al., 2006, Hwangbo et al., 2016). High-resolution genomic analyses capable of distinguishing between strains are therefore necessary. Additionally, because shotgun sequencing data often includes extracellular DNA, the presence of exogenous DNA can artificially inflate estimates of microbial diversity and genomic potential (Carini et al., 2016, Alteio et al., 2020). In recent years, single-cell genomics (SCG) has emerged as a complementary approach to shotgun metagenomics. This method bypasses the binning step inherent in shotgun analysis and enables the analysis of individual, intact cells (Gawad et al., 2016). Most of the single amplified genomes (SAGs) are derived from intact cells (Ide et al., 2022), allowing them to be treated as complete genomic representations of individual organisms. By applying single-cell genomics to specific soil components, we may be able to determine the spatial distribution of bacteria involved in N-cycling, providing insights into both their habitats and taxonomic identities.

For the application of single-cell genomics analysis to soil samples, it is essential to first disperse the soil sample and separate microbial cells from soil particles. The soil’s characteristics, especially clay and organic matter content, significantly influence the efficiency of cell extraction (El Mujtar et al., 2022). The choice of dispersion method would be crucial in single-cell analysis, as harsher dispersion conditions can increase cell recovery but may compromise cell viability (Ouyang et al., 2021, Lee et al., 2021). To detach cells from soil particles, both physical dispersion methods (e.g., blending, sonication, vortexing, shaking) and chemical dispersion methods (using ionic or non-ionic buffers) are commonly employed in combination (Williamson et al., 2011, Lindahl and Bakken, 1995, Courtois et al., 2001, Bakken, 1985). For small sample volumes or high-throughput analyses, shaking, vortexing (with or without beads), and sonication are frequently used (Khalili et al., 2019, Frossard et al., 2016). Sonication (also called ultrasonic treatment), applied at varying energy levels, has been used to partially or fully disperse soil aggregates. This method can reveal factors affecting soil aggregate stability through stepwise disaggregation (Asano and Wagai, 2014). Sonication has demonstrated a higher recovery rate for cell extraction compared to other methods and has proven effective in isolating cells (Uhlířová and Šantrůčková, 2003, Neumann et al., 2013, Frostegård et al., 1999). However, some studies have reported negative results for cell viability following sonication (Lindahl and Bakken, 1995, Khalili et al., 2019, Ellery and Schleyer, 1984). Currently, it remains uncertain which extraction technique is most suitable for SCG analysis of soil samples.

Many technical challenges remain to understand the linkage between biophysical complexity and genomic diversity within bulk soils. To this end, the current study aimed to characterize soil bacterial community using both metagenomic and SCG approaches by focusing on water-stable macroaggregates, the soil physical subunits that (i) retain the intact micro-structure and microbial habitat, and (ii) can be isolated reproducibly. We intended to fully disperse soil aggregates to maximize cell extraction because some soil microbes are localized in the interior of aggregates (e.g., N_2_O reducers, (Mitsunobu et al., 2025)) and/or difficult to detach from soil due presumably to persistent biofilm formation and inhabitation to physically-stable subunits (El Mujtar et al., 2022, Costa et al., 2018, Almås et al., 2005).

The specific objectives of this study were: 1) to establish an extraction method that allows SCG analysis on water-resistant macroaggregates, and 2) to clarify the similarity and differences in taxonomic and functional diversity of bacterial community among the individual aggregates. Using the water-stable macroaggregates from a cropland topsoil (light clay, Acrisol) that were previously used to examine physical control on the distribution of N_2_O-reducing bacteria (Mitsunobu et al., 2025), we optimized the soil dispersion method by comparing cells obtained using two soil dispersion methods (bead-vortexing method and ultrasonic method) using microscopic observation, taking into account the trade-off between soil dispersion and cell damage. Next, we compared two dispersion techniques (bead-vortexing and sonication) and evaluated the recovery of the bacterial community based on quantitative PCR and amplicon-based bacterial composition. Finally, we characterized the N-cycle and other functional aspects of single aggregates based on high-resolution genome analysis and taxonomic and functional diversity evaluation using SCG analysis data.

## 2 Materials and Methods

### 2.1 Soil sample and incubation

Soil aggregates were collected and incubated in the same manner by Mitsunobu et al. (2025). The soil aggregates were collected from the 0- to 20cm layer of an Acrisol containing bark compost from the long-term field trial plot at a research farm of the Shizuoka Prefectural Research Institute of Agriculture and Forestry in Iwata, Shizuoka (34°45’12’’N, 137°50’’33E). The average annual temperatures and annual precipitation rates were 16.8 ℃ and 1843 mm/year, respectively (Hamamatsu, 1991-2020). In the basic soil properties, bulk density was 1.2 g/cm^3^, clay was 37.7 %, silt was 25.8 %, sand was 36.5 %, total organic carbon was 3.01 %, C/N ratio was 15, and pH (H_2_O) was 7.0. We transported the soil samples to the laboratory and stored at 4 ℃ until wet sieving.

Water-stable soil macroaggregates were isolated by immersing the field-moist soil sample in autoclaved Milli-Q water on top of a 2 mm autoclaved sieve for 10 min, followed by manual vertical movement at the rate of 60 times over a distance of 1.5 cm for 2 min. We selected the aggregates of 5 - 8 mm in diameter (Table S1) and air-dried them for three days under a fume hood covered with a paper towel to prevent the contamination of the dust.

We incubated the soil aggregates with artificial soil water which contained low level of nitric acid and glucose to induce N-cycling in a chamber simulating an atmospheric environment. Using the same methods as in Mitsunobu et al. (2025), we incubated the soil aggregates with the medium simulating soil water based on Yamazaki (2004) with a slight modification. The medium contains 210 μM glucose, 350 μM KNO_3_, and a solution of 410 μM NaH_2_PO_4_・2H_2_O, 160 μM Na_2_HPO_4_・ 12H_2_O, 84 μM MgSO_4_・7H_2_O, 200 μM CaCl_2_・2H_2_O which adjusted to pH 6.4. All media were autoclaved. We placed the soil aggregate on the autoclaved glass filter paper in a Petri dish and then dropped 1-1.5 mL of the medium per an aggregate onto the paper, which allowed it to soak into the soil aggregate a little at a time without breaking the aggregate. The incubation was carried out in the dark, at 25 °C and 48 h. We changed the air four times during incubation to simulate atmospheric conditions.

The weight of incubated aggregates was measured before and after the incubation. Aggregate diameter and volume were measured by the aggregate images picture taken from directly above using Image J tool. The outline of the aggregate was extracted from the image taken from directly above, and the Feret diameter was used as the diameter. We then calculated the aggregate volume from the diameter. Using non-incubated water-stable macroaggregates (n = 5), aggregate pore characteristics were assessed using X-ray micro-computed tomography (µCT). The µCT images of three of the five aggregates were obtained from previous study (Mitsunobu et al., 2025) which gives detail method description. Briefly, the aggregates were scanned at beamline 20B2 of SPring-8 with a resolution of 2.71–2.72 µm per voxel. The µCT images were segmented using grey scale values to distinguish pores from the solid phase. Porosity was calculated as the ratio of pore volume to total aggregate volume. The pore depth distribution was analyzed by dividing each aggregate into three regions: 0– 600 µm from the surface, 600–1200 µm, and 1200 µm to the center.

### 2.2 Cell extraction method development

#### 2.2.1 Pilot1. Conventional method

First, we tested extraction method used in previous studies that applied SCG to soils, with a slight modification (Hosokawa et al., 2022, Nishikawa et al., 2022). We first mixed 1 g bulk soil samples and a buffer (soil: buffer ratio = 1 g: 3 mL) in a 15 mL tube. The buffer we used in this study was 20 mM potassium phosphate (KPi, pH 7.5) with 0.05% Tween 80. The mixture was dispersed using a mechanical shaking technique at 120 rpm for 10 min, then allowed to stand at room temperature for 5 min. 2 mL of the supernatant was collected in 1.5 mL tube and then centrifuged at 8000×g for 5 min using a benchtop centrifuge (75004251, Thermo Fisher Scientific). These steps (collection and centrifugation) were repeated twice for purification. After the last mixture was centrifuged at 300×g for 5 min, the supernatant was collected. The supernatant was stained with fluorescent dyes, LIVE/DEAD^TM^ BacLightTM Bacterial Viability Kit (Thermo Fisher SCIENTIFIC, USA) to estimate cell numbe. Eight μL of the supernatant was mixed with 1 μL of SYTO9 (50 μM) and 1μL of propidium iodide (PI, 0.5 mg/ μL), incubated for 10 min in the dark at room temperature, and observed under a fluorescence microscopy (Eclipse Ni-U, Nikon, Tokyo, Japan).

#### 2.2.2 Pilot2. Comparison of two aggregate dispersion techniques (bead-vortexing and ultrasonication)

We compared the following two dispersion treatments using 0.5 and 0.25 g of bulk soil (dry weight). In the bead-vortexing treatment, the incubated soil was dispersed in KPi buffer (soil: buffer ratio = 1 g :3 mL) with sterile zirconia beads (3 mm in diameter, soil: beads ratio = 1 g: approximately 3 g, equivalent to 16 beads). The dispersion was performed by vortexing for 1 min using a small vortex mixer (N-81, Nissin, Japan). For the 0.25 g treatment, vortexing was conducted for 0.5 min, according to Whiteley et al. (2003) and Griffiths et al. (2003). For the sonication treatment, the soil plus buffer mixture prepared in the same way as for the bead-vortexing treatment was dispersed using an ultrasonic homogenizer VP-300 (Taitec, Japan) at a low energy intensity of 30 J/ mL (15% intensity cycle of 20 s on, 20 s off), according to Neumann et al. (2013). The tip of the probe (VP-MT03) was immersed 10 mm into the soil suspension and the sonication was done in an ice bath to reduce cell damage. The mixture after each dispersion treatment was centrifuged at 300×g for 5 min, and the supernatant was collected.

The degree of aggregate dispersion was assessed by comparing the particle size distribution of single aggregates after the three dispersion treatments (mechanical shaking, bead-vortexing and sonication) by laser scattering particle size distribution analyzer ((LA-920, HORIBA Corporation, Japan). Three replicates (aggregates) of each treatment were performed. In other words, the volume of particles included in the distribution model on the small size side (< 1.98 μm) was compared. This threshold was identified as the volume-based local minimum between the two modal peaks, located in the 1.73 -1.98 μm particle size class range in the sonication and bead-vortexing dispersion samples.

We compared the cell number on LIVE/ DEAD staining in each dispersion treatment as in Pilot1. We observed large amounts of fine soil particles especially after the sonication dispersion (Fig. S1). We thus explored purification techniques to remove the soil particles from the suspension.

#### 2.2.3 Pilot3 : Purification technique after dispersion

We firstly tested the density gradient method using Nycodenz (Kallmeyer et al., 2008). The supernatant was collected from 10 g and 0.5 g incubated soil after the sonication treatment. The Nycodenz (Serumwerk Bernburg AG, Bernburg, Germany) with 1.3 g/ml density (60% w/v) was carefully layered under the supernatant in a ratio of supernatant: Nycodenz = 2:7. The sample tube were centrifuged at 10000× g for 20 min, resulting in a sharp band of bacterial cells. Then we carefully collected that to a new tube, then moved to microscopic observation. For 10 g soil, more than 10^8 – 9^ cells / soil sample was observed. For 0.5 g soil, however, the sharp band of bacterial cells was not observed after density separation. We thus concluded that Nycodenz purification was not suitable for single aggregates.

Next, we tested the sequential washing technique with 0.5 g and 0.25g of bulk soil, following Bakken (1985). The ca 500 μL and 250 μL of supernatant (S0) was transferred to new tubes after each dispersion treatments described above and centrifuged at 600×g for 15 min, resulting in the first supernatant (S1). After S1 was removed, 500 μL KPi buffer was added to the residue (R1) and subjected to repeated mixing-centrifugation steps, resulting in a series of supernatants (S2 and S3) and RS3. We observed the degree of dispersion and the cell number of each supernatant (S1, S2 and S3 in Figs. 1 and S1).

**Fig. 1.**
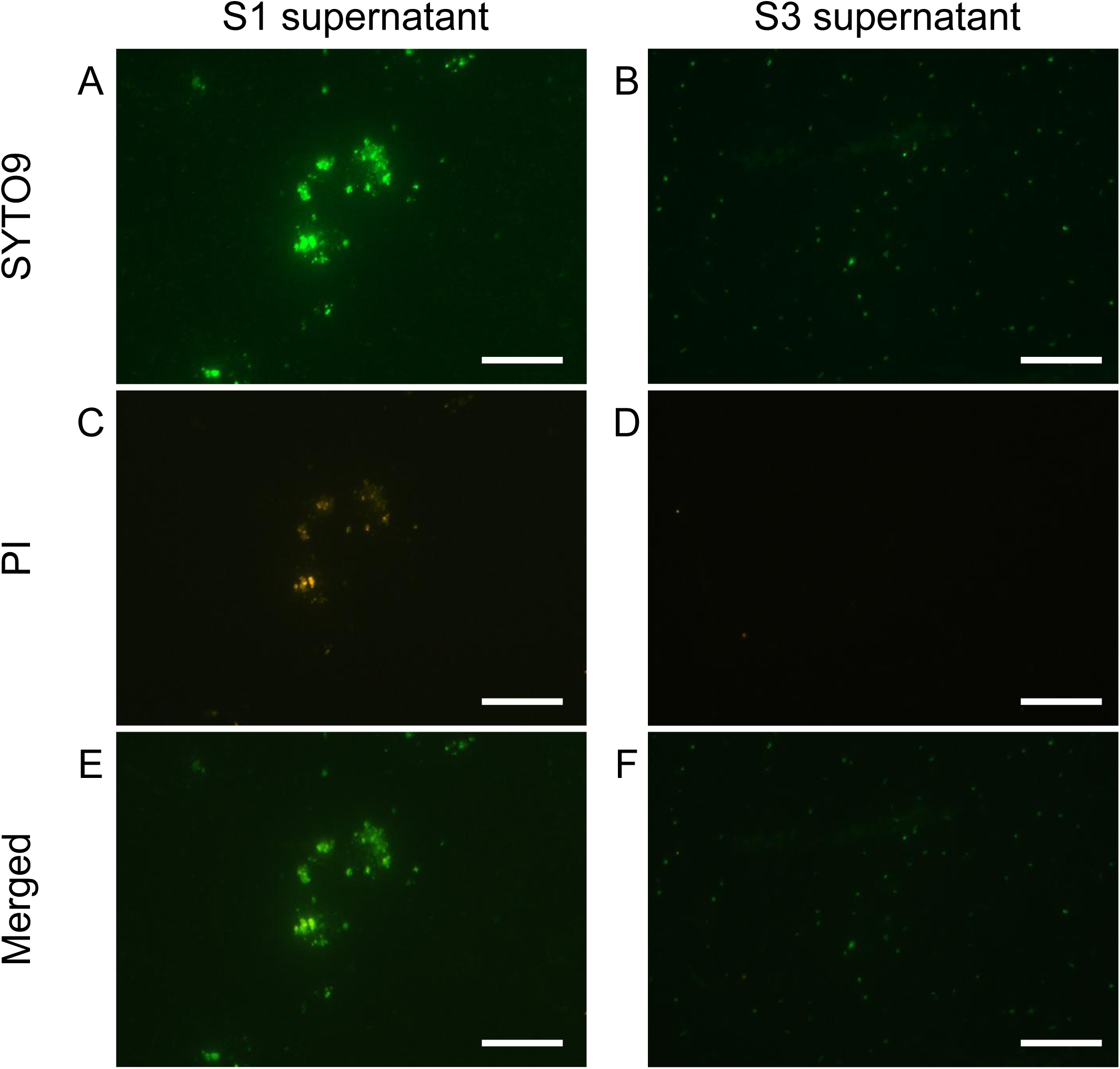
Live/dead staining of thesoilsupernatants obtained from repeated washing steps with sonication and dispersion. (A, C, E) 1st supernatant (S1) and (B, D, F) 3rd supernatant (S3) were stained with SYTO9 and PI. SYTO9 (A, B), PI (C, D) and merged (E, F) images are shown. The bar indicates 50μm. S3 has fewer PI-stained particles, indicating that it was purified to cells only.

### 2.3 Final cell extraction method (Fig. 2)

For the bead-vortexing treatment, sterile zirconia beads (3 mm-diameter) in each 1.5 mL tube and vortexed for 1 min. For the sonication treatment, an incubated aggregate was filled up with KPi buffer to 4 mL in 15 mL centrifuge tube. The mixture was dispersed using an ultrasonic homogenizer (30 J/ mL, 4W, cycle of 20 s on, 20 s off) cooling on ice. In case that a soil aggregate was above 0.1 g (dry weight), filled up to 8 mL. The mixture treated with each dispersion technique were centrifuged at 300×g for 5 min. The ca 200 μL in the bead-vortexing treatment or 3.8 mL in the sonication treatment of supernatant (S0) then was collected to 1.5 mL tube. The first supernatant (S1) was removed, and the residue (R1) and new 100μL KPi buffer were subjected to repeated mixing-centrifugation steps with same manner as Pilot3. We proceeded the third supernatant (S3) to the following steps.

### 2.4 Single-cell genomic analysis

We used total of six aggregates (three per dispersion treatment) for single-cell genomics. SAGs of soil microbes were obtained using the SAG-gel method (Chijiiwa et al., 2020, Nishikawa et al., 2022) from the third supernatant (S3) resulting from two dispersal techniques and purification (2.3). The concentration of cells was determined using LIVE/DEAD BacLight bacterial viability assay (Thermo Fisher Scientific), and the cells were then suspended in DPBS with 1.5% low-gelling-temperature agarose (Sigma-Aldrich, MO, United States) at 1 cell/capsule. After microfluidic single-cell encapsulation in the capsules, the single-cell-encapsulating gel capsules were recovered in the aqueous phase. Then, gel capsules were immersed in Buffer D2 to denature DNA, and multiple displacement amplification was performed for 3 h using the REPLI-g Single Cell Kit (QIAGEN, Hilden, Germany). After multiple displacement amplification, gel capsules were stained with SYBR Green (Thermo Fisher Scientific). FACSMelody cell sorter (Becton, Dickinson and Company, NJ, United States) equipped with a 488 nm excitation laser was used to sort the gel capsules with confirmed DNA amplification into 384-well plates at 1 bead/ well. Following capsule sorting, the 384-well plates were stored at −20°C. For the sequencing analysis, SAG libraries were prepared from the capsule-sorted plates using the xGen DNA Lib prep EZ UNI (Integrated DNA Technologies, Inc., IA, United States). Ligation adaptors were modified to TruSeqCompatible Full-length Adapters UDI (Integrated DNA Technologies, Inc., IA, United States). Each SAG library was sequenced using the DNBSEQ-G400 2 × 150 bp configuration (MGITech CO., Ltd., Beijing, China) with the MGIEasy Universal Library Conversion Kit.

Adapter sequences and low-quality reads were eliminated from raw sequence reads of single-cell genome sequences using bbduk.sh (version 38.90) with following options (ktrim=r ref=adapters k=23 mink=11 hdist=1 tpe tbo qtrim=r trimq=10 minlength=40 maxns=1 minavgquality=15).The reads mapped to the masked human genome were eliminated using bbmap.sh (version 38.90) with following options (quickmatch fast untrim minid=0.95 maxindel=3 bwr=0.16 bw=12 minhits=2 path=human_masked_index qtrim=rl trimq=10). The data of human masked index, hg19_main_mask_ribo_animal_allplant_allfungus.fa.gz, was obtained from https://zenodo.org/record/1208052#.X1hBFWf7SdY. These quality-controlled reads of single-cell genomes were assembled de novo into contigs using SPAdes (v3.15.2) (Bankevich et al., 2012) with the following options (--sc --careful --disable-rr --disable-gzip-output –k 21,33,55,77,99,127).

Contigs shorter than 200 bp were excluded from the SAG assemblies. CDSs, rRNAs, and tRNAs were predicted from the SAGs using Prokka (v1.14.5) (Seemann, 2014). with the following options (--rawproduct --mincontiglen 200). The quality of the contigs was evaluated using QUAST (v5.0.2) with the default options. The completeness and contamination of SAGs were evaluated using CheckM v1.1.3 (Parks et al., 2015) in lineage workflow with the options (-r --nt) or in taxonomy workflow with the options (--nt domain Bacteria). Taxonomy identification was performed using GTDB-Tk v2.1.0 (Chaumeil et al., 2022) with default options, and GTDB release 207. The quality of each SAG was determined based on Bowers et al. (2017). SAGs that were < 50% estimated completeness were considered low quality SAGs. SAGs that had ≥ 50% estimated completeness and < 10% estimated contamination were considered to be at least medium quality. To determine if a SAG was high quality, in addition to having > 90% estimated completeness and <5% estimated contamination, SAGs need to have 23 S, 16 S, and 5 S rRNA genes and at least 18 tRNAs present in the final assembly.

The coding sequence (CDS) and amino acid sequence of each SAG was deduced with Bakta v1.4.2 (Schwengers et al., 2021) annotation pipeline, and annotations of each CDS, were determined using eggNOG-mapper v2.1.9 (Cantalapiedra et al., 2021) with eggNOG DB version: 5.0.2 (Huerta-Cepas et al., 2019). For the SAGs of Psudoarthrobacter 87 strains that matched each other in almost full length of 16S rRNA, the gene structures of N-cycling related genes were compared and visualized using GenomeMatcher. The presence or absence of the following genes related to extracellular polysaccharide production (Cania et al., 2019) was then checked based on the KEGG IDs The percentage of SAGs with at least one detected for each exopolysaccharides (EPS) gene was calculated, and the significance difference was tested using Student’s t-test with a threshold of 0.05. Genes encoding enzymes relevant for N-cycling were detected by diamond (Buchfink et al., 2015) blastp search of all CDSs against NCycDB (Tu et al., 2019), a curated database of N-cycling genes, with the threshold of both identity and query coverage greater than or equal to 70%. To evaluate the pathway coverage in the N cycle (Tu et al., 2017), the abundance of functional genes annotated by NCycDB were compared between reaction paths and between aggregates. The sample number of SAGs for each aggregate was 226, 241, 269 and 206, 227, 203 SAGs, for Beads # 1-3 and Sonic # 1-3, respectively. With respect to SAGs with *nosZ*-like genes, we examined whether *nos* accessary genes locate adjacent to the *nosZ* gene by using interProScan 5.65-97.0 (Jones et al., 2014) in EMBL-EBI with default parameters to detect protein signatures of *nosDFLY*. The structures of *nos* gene cluster of SAGs were compared with reference genomes of related isolates or MAGs by using GenomeMatcher v3.04 (Ohtsubo et al., 2008).

### 2.5 Metagenomic analysis

We compared the supernatants and the soil residues to assess the difference in cell number and bacteria community composition to assess whether the extracted cells were representative of the soil aggregate. DNA was extracted from a total of 11 aggregate-derived R0 (0.085-0.302 g) and S2 (0.062 – 0.232 g) using an Extrap Soil DNA Kit Plus ver. 2 (BioDynamics Laboratory Inc., Tokyo, Japan) according to the manufacturer’s instructions. These 11 aggregates contained the 4 aggregates analyzed for SCG.

16S rRNA, *nosZ*-I and -*nosZ*-II amplicon sequencing and subsequent bioinformatics analysis are the same as the previous studies (Bamba et al., 2024, Hara et al., 2024). For 16S rRNA amplicon, taxonomy was assigned to ASVs using the SINTAX algorithm (Edgar, 2016) implemented in USEARCH (v11.0.667) against the SILVA database v123 (Quast et al., 2013). The sequence reads of each sample were rarefied to 27,326 reads per sample. For *nosZ* amplicon, to according to previous study (Mitsunobu et al., 2025), the generated OTU sequences were subjected to Diamond blastx against the database of NosZ amino acid sequences retrieved from the Fungene *nosZ* repository (Fish et al., 2013). Class level analysis was done based on NCBI taxonomy. The sequence reads of each sample were rarefied to 282 reads in *nosZ*-I and 856 reads per sample in *nosZ*-II.

### 2.6 Quantitative PCR analysis

Quantitative PCR (qPCR) assays were conducted using the fluorescent dye SYBR Green (THUNDERBIRD Next SYBR qPCR mix, TOYOBO) by a QuantStudio 3.0 real-time PCR System (Applied Biosystems/Thermo Fisher Scientific). 16S rRNA and genes coding for the two known clades of N_2_O reductase, *nosZ*-I and *nosZ*-II, were quantified using the primer pairs Bact1369F/ProK1492R (Suzuki et al., 2000) for the V3-V4 region of the 16S rRNA, nosZ-F/nosZ-R (Rich et al., 2003) for *nosZ*-I and nosZ-II-F/nosZ-II-R (Jones et al., 2013) for *nosZ*-II. The PCR reactions for 16S rRNA started with an initial denaturing step of 95 °C for 30 s, followed 40 cycles at 95 °C for 5 s and 60 °C for 30 s. Thermal cycling conditions for *nosZ*-I were initial denaturation at 95 °C for 30 s, 40 cycles at 95 °C for 5 s, 58 °C for 30 s, 72 °C for 30 s. For *nosZ*-II, annealing temperature was 54 °C. Melting curve analyses involved a denaturing step at 95 °C for 15 s, annealing at 65 °C for 1 min and melting in 0.1 °C steps up to 95 °C. Standard curves for each assay were generated by serial dilutions of linearized plasmids with cloned fragments of environmental DNA. Amplification efficiencies were 101.7% for 16S rRNA, 110.8% for *nosZ*-I and 84.2% for *nosZ*-II.

### 2.7 Data analysis

For alpha diversity analysis, observed ASVs or OTUs, Chao1, Shannon, InvSimpson and Faith’s phylogenetic diversity index were calculated for amplicon and SCG analysis sample. Rarefaction interpolation and extrapolation analysis of taxonomic richness (the observed ASVs) and functional richness (number of KEGG-ID and number of different kinds of N-cycling related genes annotated by NCycDB) was performed using iNEXT v 3.0.0.packages (Hsieh et al., 2016). Furthermore, we investigated the shape of the relationship between species (OTUs based on 16S rRNA) and functional (KEGG-ID) richness to compare redundancy patterns within individual aggregate (Gamfeldt et al., 2008, Teichert et al., 2017). This relationship is expected to be linear when all species of an assemblage support singular functions, meaning that the loss of any species will produce an important and equivalent decline in functional richness. On the contrary, functionally redundant assemblages will display curvilinear relationships, i.e. saturation trends, as some functional traits are shared by multiple species.

For beta diversity analysis, a weighted UniFrac distance matrix was employed in a principal coordinate analysis (PCoA). Subsequently, a permutational multivariate analysis of variance (PERMANOVA) was conducted. The linear discriminant analysis effect size (LEfSe) method determines the features (clades, operational taxonomic units) most likely to explain differences between classes by combining standard tests of statistical significance with additional tests that encode biological consistency and effect relevance (Segata et al., 2011). Soil unit bioindicators at the phylum level were determined using LEfSe analysis with a p-value of 0.05.

For molecular phylogenetic analysis focused on Gemmatimonadota, as the most abundant in soil environments and potentially important role in reducing the N_2_O (Chee-Sanford et al., 2019, Jones et al., 2013, Oshiki et al., 2022), the alignment of nucleic acid was performed using MAFFT (Katoh et al., 2019). A phylogenetic tree was constructed in MEGA 7 (Kumar et al., 2016) using the maximum likelihood method by best fit model, (Tamura-Nei model for 16S rRNA, General Time Reversible model for *nosZ*) selected best-fit model (Tamura et al., 2011) in bootstrap analyses based on 500 replicates.

All statistical analyses in community ecological analysis were performed in R software v4.2.1 using phyloseq v1.42.0 (McMurdie and Holmes, 2013), ggplot2 v3.5.0 (Wickham, 2011), microeco v1.4.0 (Liu et al., 2021), vegan v2.6-4 (Oksanen et al., 2013), cowplot v1.1.3 (Wilke, 2015). and ape v5.7-1 (Paradis and Schliep, 2019).

## 3 Results

In this study, we first showed that the combination of dispersion (bead-vortexing or ultrasonic) and sequential washing as the feasible method to extract intact cells from the soil aggregates for SCG analysis. Second, we performed SCG analysis using this method and showed the difference between the two dispersion techniques in their extractability of bacterial cells and DNA with respect to genome quality as well as their taxonomic and functional diversity. Third, by analyzing selected functional genes related to exopolysaccharides (EPS) and N-cycling, we also showed the two dispersion techniques resulted in important differences in the recovery of these genes.

### 3.1 Development and evaluation of soil bacterial extraction method

#### 3.1.1 Comparison of protocols for soil bacterial extraction

We compared previous and new methods of extracting bacterial cell from the soil aggregates for subsequent SCG analysis. Pilot1, using the previous extraction method (Hosokawa et al., 2022, Nishikawa et al., 2022), resulted in insufficient cell extraction for the studied soil (Fig. S1A). Next, we compared the two dispersion techniques (bead-vortexing and sonication treatment, Pilot2). The post-dispersion soil suspension had a bimodal particle size distribution with greater liberation of smaller-sized particles (ca. 0.5 µm) at the expense of larger-sized particles (ca. 15-90 µm) in the sonication treatment compared to the bead-vortexing treatment. In contrast, the suspension of mechanical shaking showed clearly incomplete dispersion – much greater volume of >50 µm particles and much less volume of < 1 µm particles compared to the other two dispersion techniques (Fig. S2). Specifically, the total volume of particles smaller than the bimodal distribution boundary value of 1.98 µm was 54.5 ± 1.6% after the sonication and 45.4 ± 9.0% after the bead-vortexing treatments (t-test, p = 0.1), which confirmed a greater degree of the aggregate dispersion by the sonication treatment in agreement with our visual inspection.

Aggregate pore characterization by X-ray µCT showed that the majority of pore was present in outer zones and the interior zone (deeper than 1.2 mm from the aggregate surface to the center) accounted for only 0-13 % of total pore volumes (Fig. S3) among the five randomly-picked aggregates (diameter range: 4.9-7.1 mm). The pore distribution pattern suggest that outer zones are less physically stable and thus more susceptible for dispersion than the interior zones of these aggregates.

Cell counts after the two dispersion treatments was >10^6^ (cell/ sample), which was above the minimum counts (10^6^ cell/ sample) for SCG analysis (Fig. S1B, C). In the extracted solution, we, however, observed many suspended soil particles (particularly after the sonication treatment) that would interfere with SCG analysis.

We thus conducted a sequential washing after the dispersion treatments to remove the suspended particles that would interfere SCG analysis (Pilot3). The supernatants recovered after the first (Fig. 1A, C, E) and second washings (data not shown) still contained some soil particles (microaggregates) as well as bacterial cells even after the sonication treatment. In the supernatant after the third washing, we hardly observed any aggregated particles and, based on nucleic acid staining, the number of non-bacterial particles was reduced (Fig. 1E, F) while achieving the bacteria count of 10^6-7^ per sample (Fig. S1E, F). Based on these results, we adopted the soil extraction method which combines the aggregate dispersion with the washing technique for the aggregate-scale SCG analysis (Fig. 2).

**Fig. 2.**
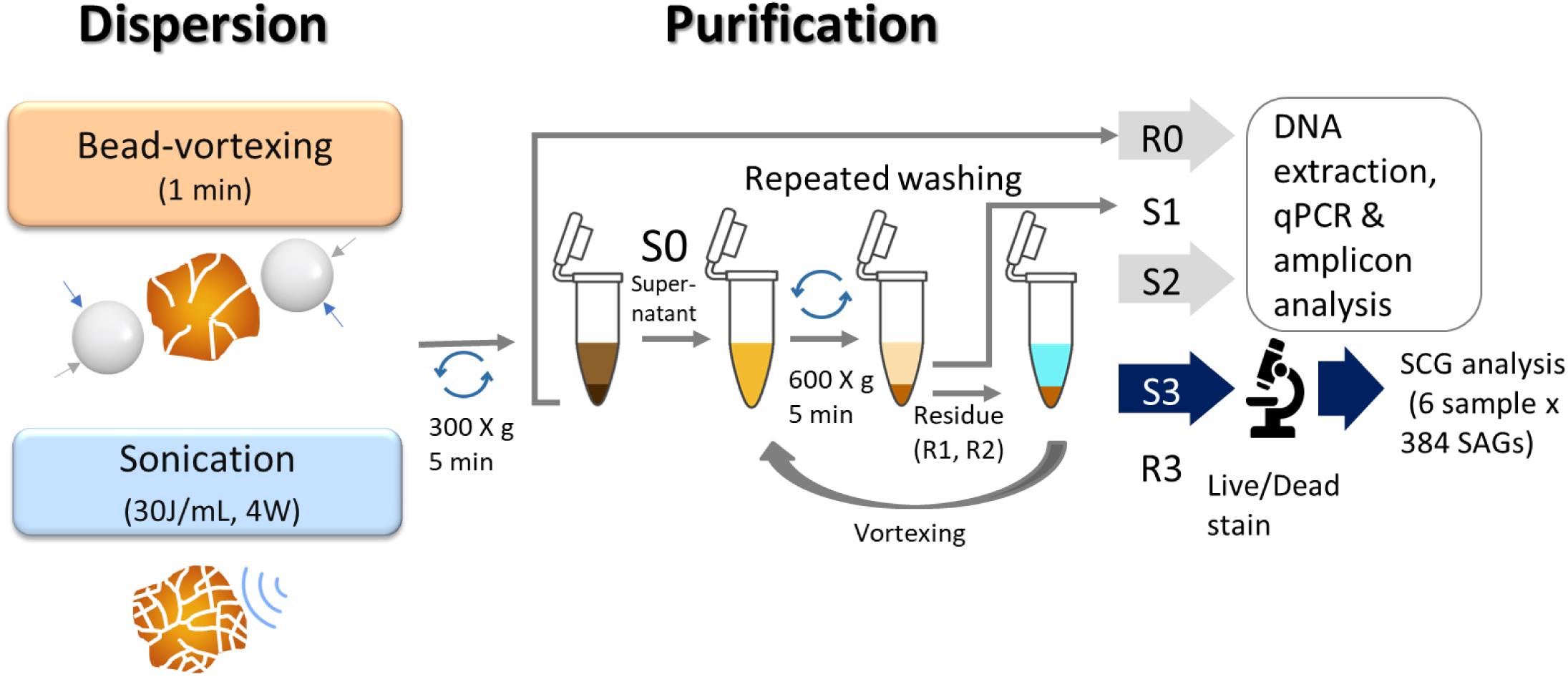
Illustration of a protocol for extraction of soil bacteria. We used R0 and S2 for qPCR and amplicon analysis and S3 for single cell genomics analysis.

#### 3.1.2 Comparison of bacteria community extracted from soil residues and supernatants

We compared the difference in copy number and bacteria community composition between the second supernatants (S2 in Fig. 2) and the soil residues after the initial centrifugation (R0 in Fig. 2) to assess the extent to which the extracted cells represent the microbiome of the single soil aggregates. The copy number in the supernatant was one to two orders of magnitude lower than that in the soil residues (Table S1). When comparing the two dispersion techniques, the copy number of 16S rRNA, *nosZ*-I, and *nosZ*-Ⅱ tended to be higher in the sonication compared to the bead-vortexing treatment while the significant difference was detected only for *nosZ*-Ⅱ (Fig. 3C, t-test, p= 0.02).

**Fig 3.**
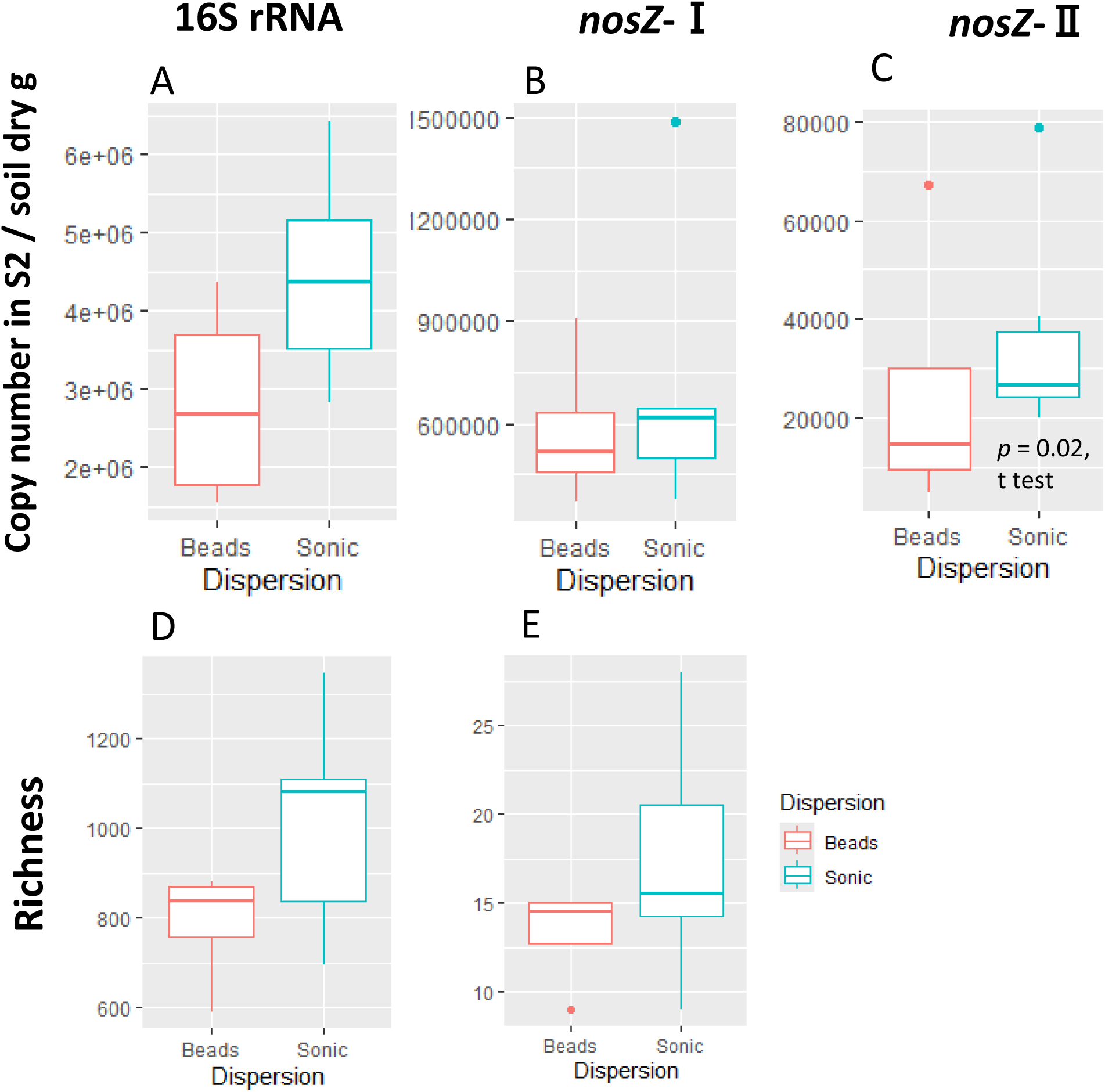
Copy number and ASV/OTU richness of the second supernatant (S2). The copy number for 16S rRNA(a), nosZ-I(b) and -Ⅱ (c) and the richness (observed ASV/OTU)for 16S rRNA(d)and *nosZ*–I(e). We didn’t perform amplicon analysis for *nosZ*–Ⅱ regiondueto not enough sample volume. The significant difference were determined by t-test after outlier exclusion.

The amplicon analysis of individual aggregates showed that S2/R0 ratio of ASV for 16S rRNA was 0.43 on average (range: 0.31 – 0.57), that for *nosZ*-I OTU was 0.24 (0.10 – 0.41) (Table S1). On average in each aggregate, 53% (range to 30-70%) and 34% (14 -57%) of ASVs for 16S rRNA and OTUs for *nosZ*-I in S2 were shared between R0 and S2 of total ASVs or OTUs. The Shannon diversity index of the S2 community (16S rRNA) was higher by in the sonication (5.68 ± 0.29) compared to the breads treatment (4.70 ± 0.17) (Table S1, t-test, p=0.01).

The difference in bacterial community composition between the supernatant and the residue as well as between the two dispersion treatments was shown by PCoA of the weighted UniFrac distance based on16S rRNA region (PERMANOVA, Fraction: r^2^ = 0.40, F =14.8, p = 0.001, Dispersion: r^2^ = 0.065, F =2.4, p = 0.047, Fig. 4A). The distance between S2 and R0 in the sonication treatment (0.31 ± 0.02) was significantly smaller (t-test, p < 0.001) than that in the bead-vortexing treatment (0.36 ± 0.03). In contrast to 16S rRNA, no significant difference between the two soil fractions and two dispersion treatments was found for *nosZ*-I and -Ⅱ regions (Fig. S4A, C). We noted that *Bradyrhizobium* in *nosZ*-I and *Cloacibacterium* and *Runella* (both Bacteroidota) in *nosZ*-Ⅱ appeared to be higher in the sonication compared to the bead-vortexing treatment (Fig. S4B, D). The characteristic phylum in S2 shown by the LEfSe analysis was Actinobacteriota, Parcubacteria for the bead-vortexing treatment and Armatimonadota, Bacteroidota, Candidatus_Saccharibacteria and Gemmatimonadota for the sonication treatment (Table S2).

**Fig 4.**
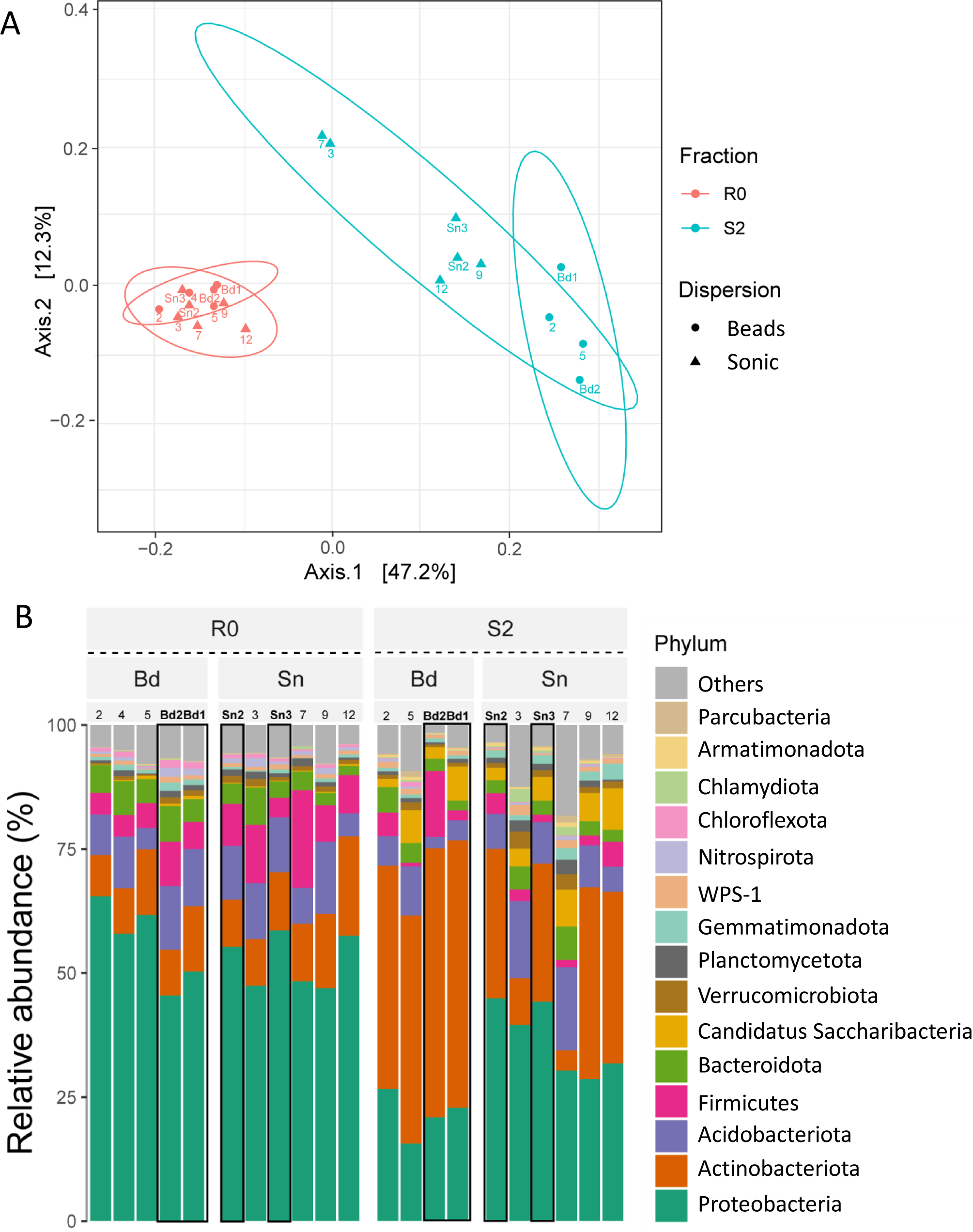
Community differences are shown by a PCoA plot of the weighted UniFrac distance matrix (a) and a bar chart at the phylum level (b) across the different fractions and dispersion. R0, residue; S2, second supernatant; Bd,after beads treatment; Sn, after the sonication treatment. The numbers indicate aggregate sample number. Sn2-3 and Bd2-3 Sn3 highlighted in bold with black border, indicate the Sonic and Beads treatment sample in the SCG analysis.

### 3.2 Profiles of single amplified genomes extracted from single aggregates

#### 3.2.1 SAGs data quality

We evaluated basic genomic information including its quality based on the 2,304 SAGs analyzed. Of all SAGs, 2 SAGs were classified as high-quality (HQ), 241 SAGs as medium-quality (MQ), and 2047 SAGs as low-quality (LQ) genomes (Table 1). The remaining 14 SAGs had a contamination rate of 10-34 %, and most of them were Actinobacteriota. The average total length was 3.72, 2.52, and 0.91 Mb, the average N50 was 101, 60, and 25 Kb, and the average number of tRNAs were 39.5, 27.9, and 10.6, for HQ, MQ, LQ, respectively. When comparing the two dispersion techniques, SAGs of HQ and MQ tended to be higher in the bead-vortexing treatment than in the sonication treatment with an average of 15.5 % and 5.4 %, respectively. Similar trends were also observed for the completeness, the number of tRNA and total CDS while no significant difference was shown for N50 (Table 1, Fig. S5). When analyzing after excluding the most dominant Actinobacteriota SAGs, all indexes tended to diverge to a greater extent between the aggregates compared to between the dispersion treatments (Fig. S5).

**Table 1.**
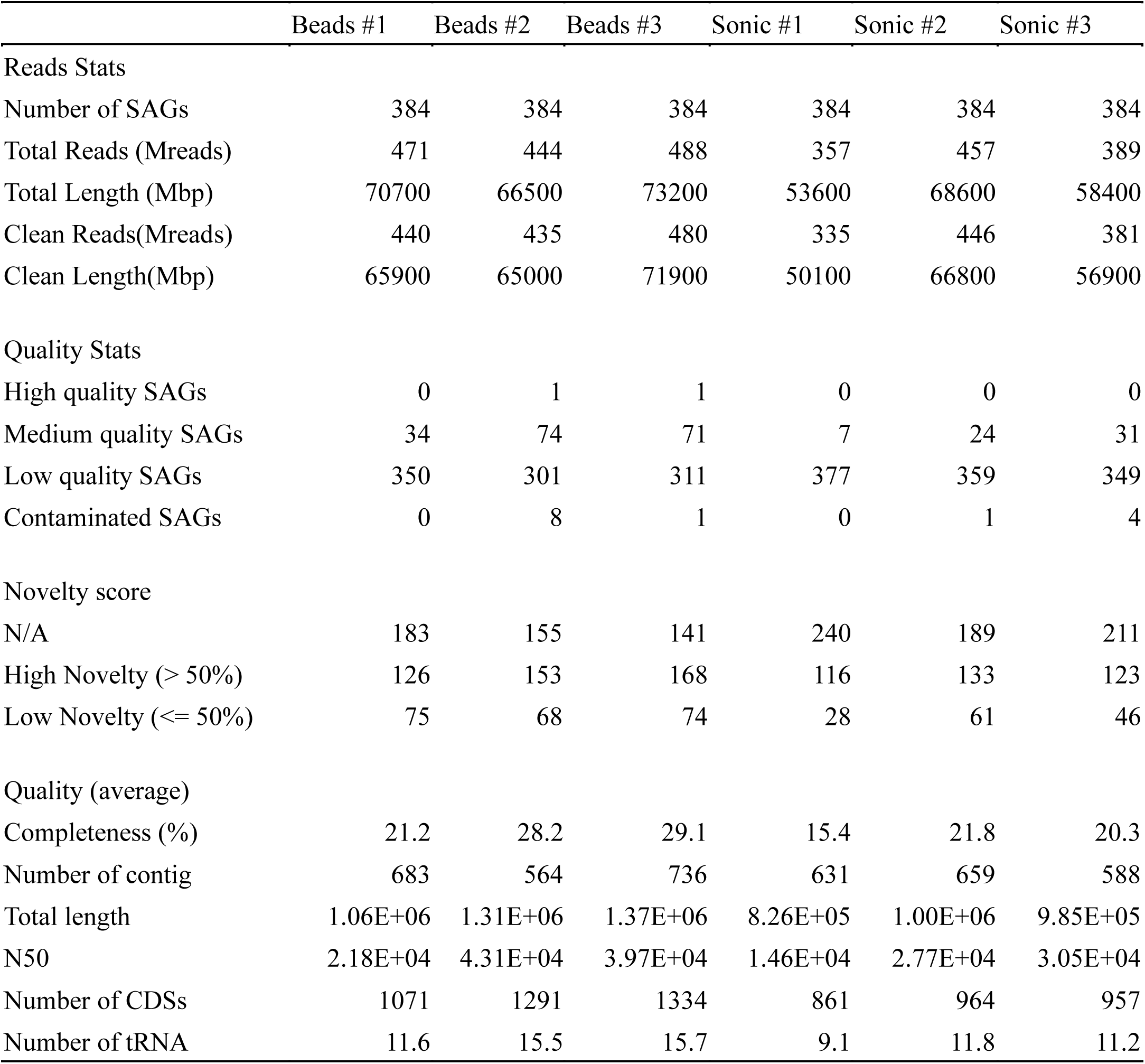
Basic genomic information and quality, which are the evaluation indices for single-cell analysis.

#### 3.2.2 Taxonomic and functional diversity

Based on phylogenetic annotation based on the GTDB, Actinobacteriota (66.0%, 1034/ 1566 SAGs identified to the phylum level) was the most frequent phyla (Fig. 5A). When comparing the two dispersion techniques among SAGs identified by phylum level, the average number of phyla detected was 12.6 in the sonication and 9.3 in the bead-vortexing treatment. The first dominant Actinobacteriota was significantly more frequent after the bead-vortexing treatment (209.0 ± 11.9 SAGs) than after the sonication (135.7 ± 16.8 SAGs) (t-test, p= 0.007). In contrast, the second dominant Proteobacteria and the third dominant Acidobacteriota tended to be higher after the sonication (Fig. 5A, Fig. S6). The results indicated that the cell extraction with the sonication dispersion allowed the detection of a wider range of phylum compared to that with the bead-vortexing dispersion technique. We calculated Faith’s phylogenetic diversity using the full-length 16S rRNA data. The phylogenetic diversity appeared to be higher after the sonication treatment (175.1 ± 15.4) compared to the bead-vortexing treatment (155.6 ± 14.7) but without statistically significant difference (t test, p =0.2). Similar trend was also observed in the rarefaction curve of ASV (Fig. 5B).

**Fig. 5.**
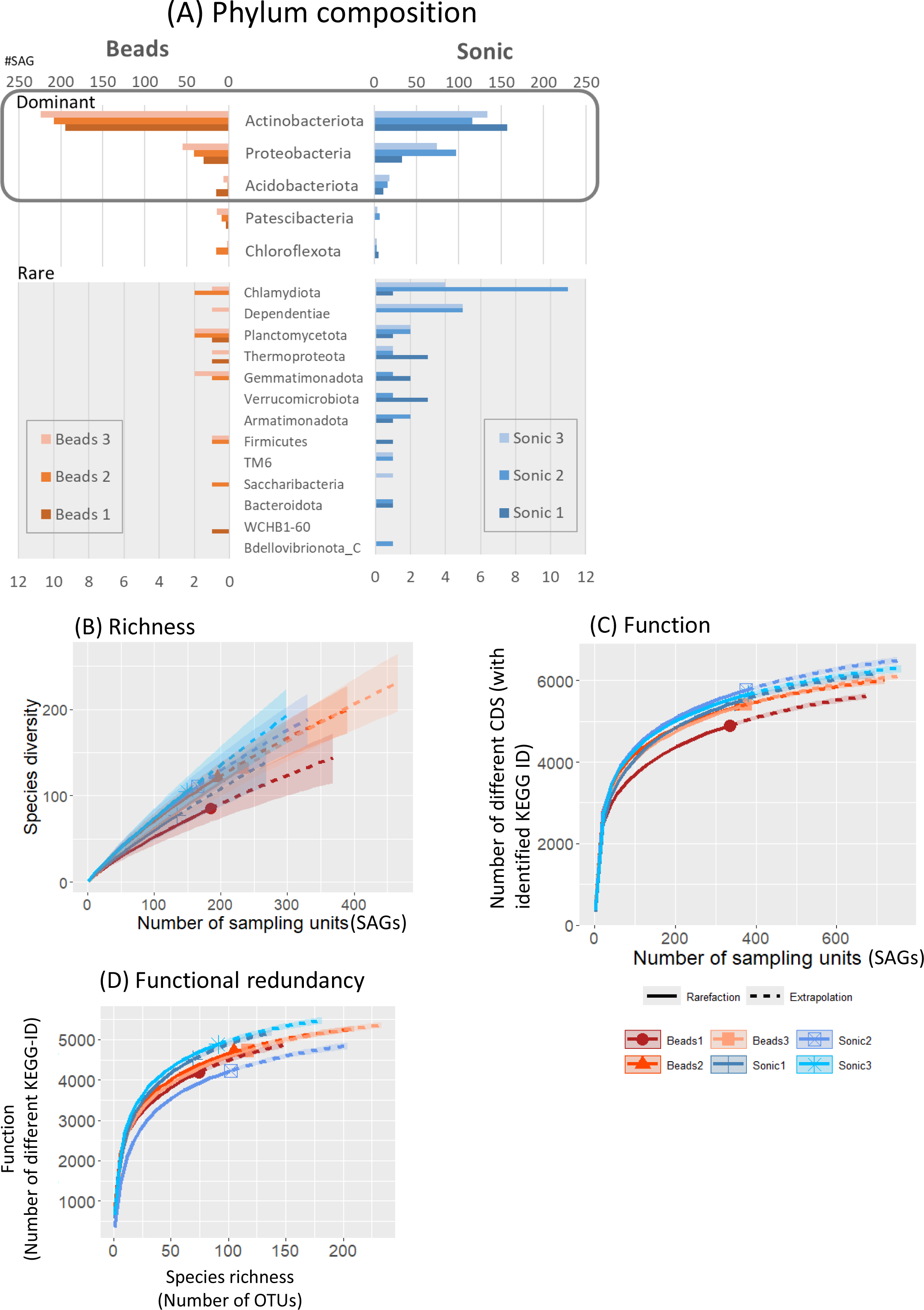
Taxonomic and functional diversity of SAGs. The taxonomic composition of SAGs, annotated based on the GTDB at the phylum level, is presented for all SAGs (A). Rarefaction curves illustrate alpha diversity, including species richness based on 16S rRNA gene sequences (B), functional diversity (C), and functional redundancy (D). The distribution of SAG counts at the genus level for the three most abundant genera within each phylum is detailed in Supplementary Figure S6.

We also assessed the functional diversity of the SAGs. When comparing the two dispersion techniques among SAGs, the number of different CDS based on KEGG-ID tended to be higher in the sonication compared to the bead-vortexing treatment (Fig. 5C). The comparison of the number of functional genes with species richness showed that total number of functional genes was equally high (4000-5000) with similar plateau shapes for all the aggregates, suggesting high bacterial functional redundancy across the six single aggregates (Fig. 5D). However, the inter-aggregate variation became noticeable when focusing on N-cycle genes (Fig. S7A), especially on specific processes (Fig. S7B-D) (see Section 3.3.2).

#### 3.2.3 Genetic variation in comparative arrangement of gene cluster within a species

One of the advantages of single-cell analysis over MAGs is its capability of detecting the differences among the genes from closely related strains We analyzed the SAGs of Psudoarthrobacter 87 strains that matched each other in almost full length of 16S rRNA, and compared the contigs in which the assimilatory nitrate reductase and nitrite reductase clusters were located at loci of Y7B10_sc-00145, Y7B8_sc-00268, Y7B8_sc-00316, and Y7B8_sc-00152 (Fig. 6A). The results showed overall very high homology, except for a loss of about 3500 bp in Y7B10_sc-00145. In Y7B10_sc-00145, a region of about 3500 bp encoding the Radical SAM domain protein gene and the prolipoprotein diacylglyceryl transferase gene is missing, while it was conserved in the other three SAGs.

**Fig.6.**
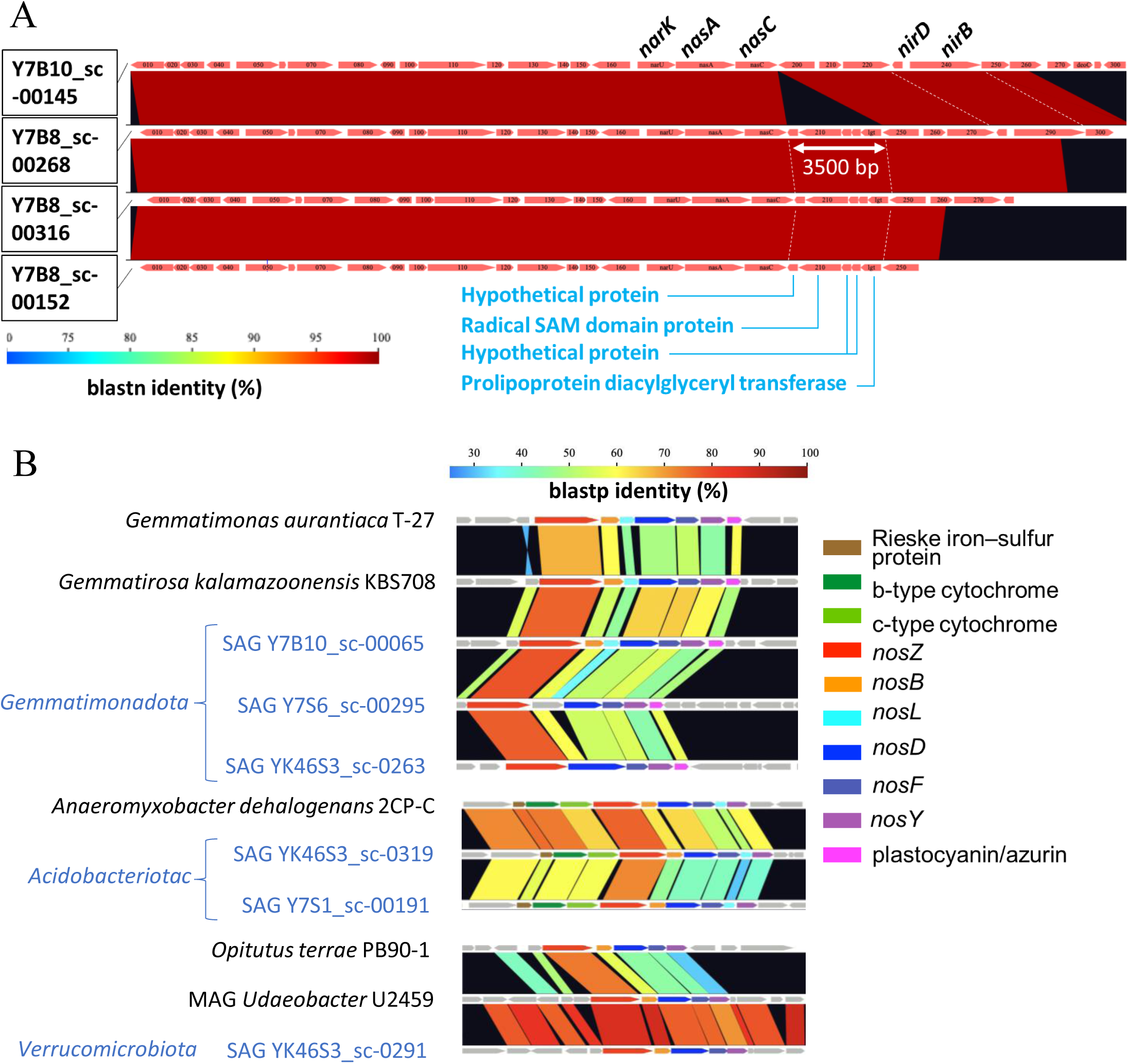
Comparative arrangement of gene cluster among SAGs. (a) Cluster for *nos* between SAGs (blue letter) and reference strains (black letter) . (b) Cluster of the assimilatory nitrate reductase and nitrite reductase cluster among four SAGs belong to Pseudoarthrobacter. Colour coding represents % homology based on nucleotide sequences of genes calculated by GenomeMatcher (Ohtsubo et al., 2008).

### 3.3 Functional gene analysis of single amplified genomes

The results above (3.2.2) showed that the dominant phyla were the same between the two dispersion treatments while the sonication treatment obtained higher functional diversity. Next, we carried out more detailed profiling on functional genes that are related to soil aggregation and N-cycling.

#### 3.3.1 Comparison of SAGs harboring EPS related genes

We investigated a total of 242 SAGs of middle and high quality for the presence of exopolysaccharides (EPS) related genes as EPS play a major role in cell attachment to soil surface and in aggregate formation (Wagai et al., 2023). A total of 242 SAGs of MQ and HQ were investigated for the presence of EPS related genes (Fig. S8). The proportion of SAGs harboring the polysaccharide export outer membrane protein gene (*wza*, KO number K0991) was significantly lower in the bead-vortexing treatment compared with the sonication treatment, 6.1% ± 3.1% and 33.7% ± 8.5%, respectively. In addition, 36 out of 61 SAGs in the sonication treatment and 16 out of 181 SAGs in the bead-vortexing treatment belonged to actinomycetes, but none possessed the *wza* gene. In contrast, many SAGs (86.2% ± 12.2% in the sonication and 72.2% ± 25.5% in the bead-vortexing treatment) in bacteria other than actinomycetes harbored the *wza* gene. The same trend was observed for LptBFGC lipopolysaccharide export complex permeasegene (*lptF*, KO no. K07091), LptBFGC lipopolysaccharide export complex permease gene (*lptG*, KO no. K11720), LptBFGC lipopolysaccharide export complex inner membrane protein gene (*lptC*, KO no. K11719), capsular polysaccharide export system permease gene (*kpsE*, KO no. K10107) (Fig. S8).

#### 3.3.2 Nitrogen cycling genes

We then narrowed down the functional analysis to N cycling genes and assessed how the frequency of major N-cycling genes CDS (annotated by NCycDB) differed among the reaction paths (arrows, Fig. 7) and among the six soil aggregates (six-cell boxes with heatmap, Fig. 7). Relatively abundant genes (> 20%, bold arrows) were *nirBD* related to NO_2_^-^→NH_4_^+^ and *narB*, *nasAB*, *narGHIJ* related to NO_3_^-^→NO_2_^-^. In addition, *nmo* and *gdh* genes that are related to organic nitrogen degradation, and a *GS* gene involved in glutamine synthesis were abundant. On the other hand, functional genes related to nitrification, nitrogen fixation and anamox were little detected. Low abundance genes (< 5%, thin arrow) were *nosZ*, *nrfA*, *norBC* and *napAB*. When comparing among the aggregates, the high abundance genes were typically found in the aggregates isolated after the bead-vortexing dispersion treatment (the upper 3 cells in each box, Fig. 7), while the low abundance genes tended to be found in the aggregates under the sonication dispersion treatment. A rarefaction estimates CDS richness tended to indicate the detection of more diverse functional genes by the sonication dispersion except for assimilatory nitrate reduction (ANR) -related genes (Fig. S7).

**Fig.7.**
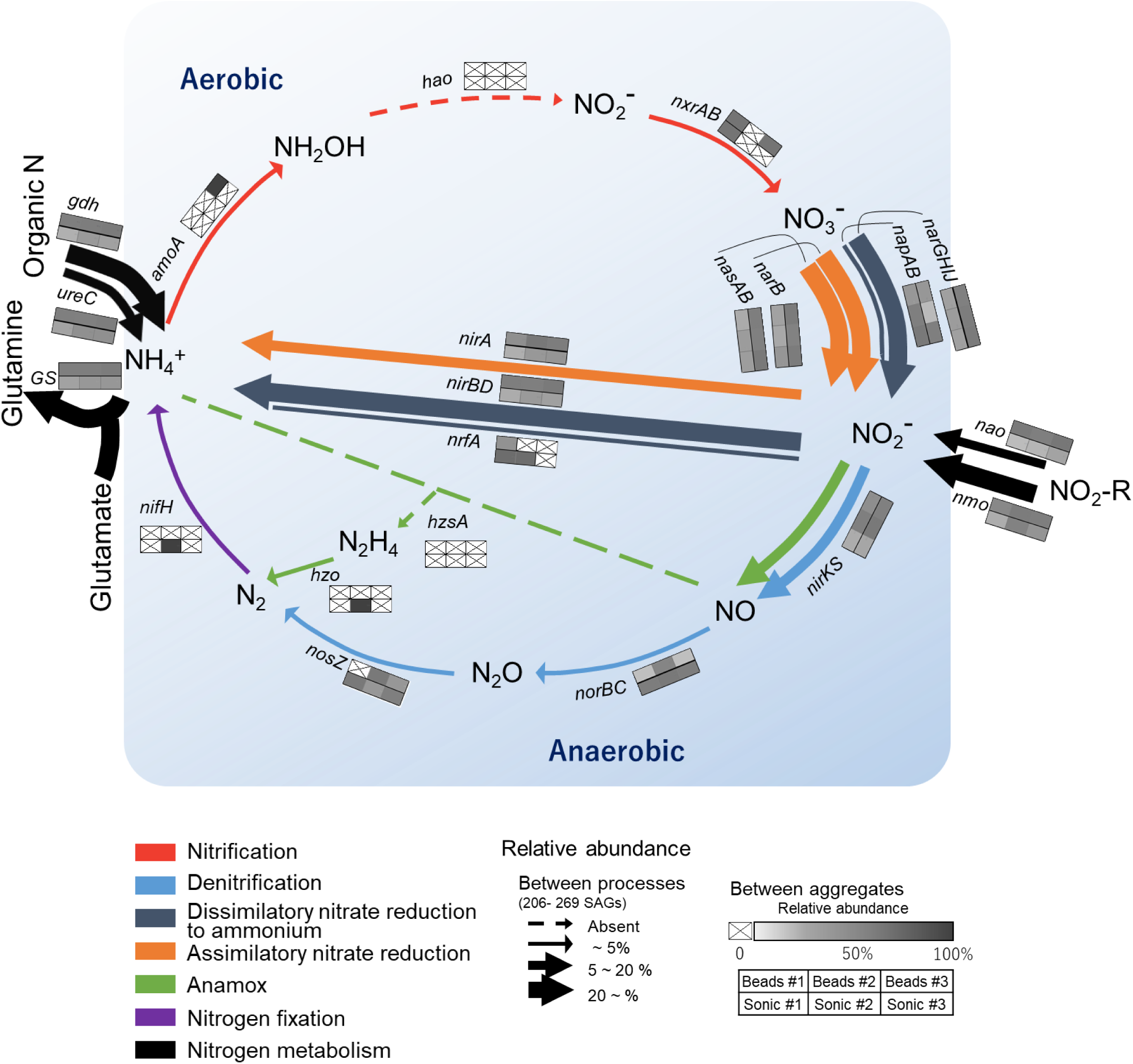
The potential coverage of pathways in the nitrogen cycle in thesingle aggregate.The sample number of SAGs for each aggregate was 226,241,269 and 206,227,203 SAGs, for Beads# 1-3 and Sonic# 1-3, respectively.

We also assessed whether each taxon (phylum) differed in the diversity of functional gene arrangements (gene arrangement/synteny) involved in the N-cycling. Actinobacteriota, the most dominant phylum, harbored multiple genes related to dissimilatory nitrate reduction to ammonium (DNRA), denitrification (nitrate and nitrite reduction), and ANR within single cells except for Thermoleophilia (class) that harbored few N-cycling genes. Proteobacteria, the second dominant phylum also harbored the genes for multiple N-cycling pathways. In contrast with Actinobacteriota, many classes of Proteobacteria harbored genes for denitrification-specific processes, such as *nirKS* and *nosZ*-I. The only SAGs with genes involved in nitrification belonged to archaea (Nitrososphaeraceae)

Homology search of CDSs of all SAGs by blastp against NcycDB identified NosZ-like CDSs from 15 SAGs (Table 2). Eleven out of these 15 SAGs were taxonomically assigned by Gtdbtk: five SAGs from Acidobacteriota, three from Proteobacteria, two from Gemmatimonadota, one from Verrucomicrobiota. Although the rest of four SAGs were unclassified by Gtdbtk, their NosZ-like CDSs showed homology to inferred NosZ of Bacteroidota, Proteobacteria, and Gemmatimonadota.

**Table 2.**
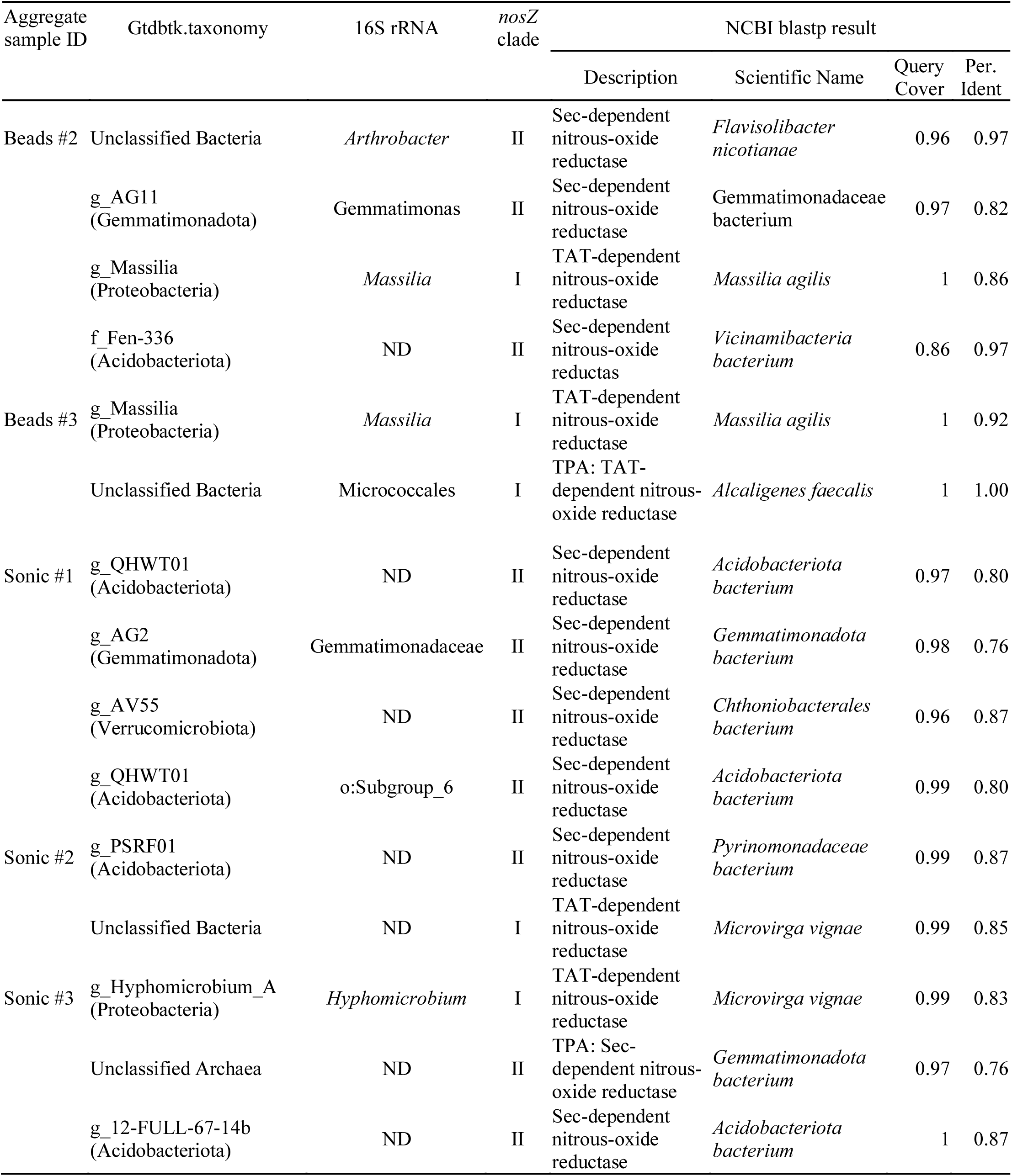
Taxonomy of *nosZ* harboring SAGs.

| Aggregate sample ID | Gtdbtk.taxonomy | 16S rRNA | *nosZ* clade | NCBI blastp result |  |  |  |
| --- | --- | --- | --- | --- | --- | --- | --- |
|  |  |  |  | Description | Scientific Name | Query Cover | Per. Ident |
| Beads #2 | Unclassified Bacteria | *Arthrobacter* | II | Sec-dependent nitrous-oxide reductase | *Flavisolibacter nicotianae* | 0.96 | 0.97 |
|  | g_AG11 (Gemmatimonadota) | Gemmatimonas | II | Sec-dependent nitrous-oxide reductase | Gemmatimonadaceae bacterium | 0.97 | 0.82 |
|  | g_Massilia (Proteobacteria) | *Massilia* | I | TAT-dependent nitrous-oxide reductase | *Massilia agilis* | 1 | 0.86 |
|  | f_Fen-336 (Acidobacteriota) | ND | II | Sec-dependent nitrous-oxide reductas | *Vicinamibacteria bacterium* | 0.86 | 0.97 |
| Beads #3 | g_Massilia (Proteobacteria) | *Massilia* | I | TAT-dependent nitrous-oxide reductase | *Massilia agilis* | 1 | 0.92 |
|  | Unclassified Bacteria | Micrococcales | I | TPA: TAT-dependent nitrous-oxide reductase | *Alcaligenes faecalis* | 1 | 1.00 |
| Sonic #1 | g_QHWT01 (Acidobacteriota) | ND | II | Sec-dependent nitrous-oxide reductase | *Acidobacteriota bacterium* | 0.97 | 0.80 |
|  | g_AG2 (Gemmatimonadota) | Gemmatimonadaceae | II | Sec-dependent nitrous-oxide reductase | *Gemmatimonadota bacterium* | 0.98 | 0.76 |
|  | g_AV55 (Verrucomicrobiota) | ND | II | Sec-dependent nitrous-oxide reductase | *Chthoniobacterales bacterium* | 0.96 | 0.87 |
|  | g_QHWT01 (Acidobacteriota) | o:Subgroup_6 | II | Sec-dependent nitrous-oxide reductase | *Acidobacteriota bacterium* | 0.99 | 0.80 |
| Sonic #2 | g_PSRF01 (Acidobacteriota) | ND | II | Sec-dependent nitrous-oxide reductase | *Pyrinomonadaceae bacterium* | 0.99 | 0.87 |
|  | Unclassified Bacteria | ND | I | TAT-dependent nitrous-oxide reductase | *Microvirga vignae* | 0.99 | 0.85 |
| Sonic #3 | g_Hyphomicrobium_A (Proteobacteria) | *Hyphomicrobium* | I | TAT-dependent nitrous-oxide reductase | *Microvirga vignae* | 0.99 | 0.83 |
|  | Unclassified Archaea | ND | II | TPA: Sec-dependent nitrous-oxide reductase | *Gemmatimonadota bacterium* | 0.97 | 0.76 |
|  | g_12-FULL-67-14b (Acidobacteriota) | ND | II | Sec-dependent nitrous-oxide reductase | *Acidobacteriota bacterium* | 1 | 0.87 |

The 16S rRNA and *nosZ* classifications did not match for two SAGs from the bead-vortexing treatment that were not unclassified by Gtdbtk. The comparing the results of the single-cell and amplicon analyses by focusing on Gemmatimonadota, revealed that the sequences of *nosZ* and of 16S rRNA from SCG were closer to the dominant ASV of Gemmatimonadota identified by the amplicon analyses of both *nosZ* and 16S rRNA than any of the known isolated strains in Gemmatimonadota (Fig. S9).

Furthermore, we examined weather nos accessory genes located in *nosZ*-flanking region of SAGs from Acidobacteriota, Gemmatimonadota and Verrucomicrobiota, the phyla known to be difficult to culture. Blastp against NCBI and InterProScan analysis detected CDSs with homology to *nosDFY*, important for formation of the active site of *nosZ*, and other nos accessory genes in their *nosZ*-flanking regions. Comparative analysis of *nosZ*-flanking regions revealed that the gene arrangement of the *nos* gene cluster was highly conserved among the SAGs and reference genomes (Fig. 6B).

## 4 Discussion

### 4.1 Optimal extraction methods for SCG and general characteristics of extracted bacteria

We were able to obtain genomic information based on SCG analysis from single soil aggregates for the first time by optimizing the extraction method after a series of pilot tests assessing aggregate dispersion techniques (zirconia bead-vortexing and sonication) as well as the purification of dispersed soil suspension. In theory, a high recovery of bacterial cells from soil is achievable by sufficient, if not complete, dispersion of the aggregates and detachment of cells from soil surfaces while minimizing the physical damage to the cells. Our results suggested that the ultrasonic dispersion was more effective than the bead-vortexing for the recovery of bacterial cells and that of particular bacterial groups (especially *nosZ* harboring bacteria in Fig. 3C). Below, we first discuss the methodological aspects in terms of the representativeness of the number and composition of bacteria recovered.

A fundamental question when applying cell or DNA extraction techniques to soil is the extent to which the bacteria extracted represent the whole microbiome in the soil sample (Pathan et al., 2021, Wydro, 2022). To a limited extent, we assessed this by comparing the extracted bacterial genes between the soil residue obtained after the soil dispersion/centrifugation (R0) and the second supernatant of the soil suspension (S2) because the third supernatant (S3) was used for the SCG analysis and thus unavailable (Fig. 2). We found that the 40% of ASVs in S2, based on 16S rRNA amplicon analysis, was common between S2 and R0 despite that the number of bacteria in the supernatant was two orders of magnitude lower than the residue (Table S1). These results suggest that the microbial community in the third supernatant used for SCG analysis is likely to be not drastically different from the community present in the bulk sample (i.e., individual macroaggregates). While the assessment of the bacteria extraction efficiency in soil is rather rare in the literature, our results are generally consistent with the previous study which examined the bacteria recovery from pasture soils in New Zealand. Highton et al. (2023) used an extraction method comparable to ours and compared between bulk soils and their water extracts. After the soil dispersion in water using a blender followed by low-speed centrifugation and washing, they used an epifluorescence microscopy based (EFM) quantification method and showed that the number of extracted cells ranged from 3.3 to 9.4% of the bulk soil. In comparison, our current study showed that the copy numbers of S2 relative to that of R0 for 16S rRNA, *nosZ*-I and -II were 5.2 %, 4.1%, and 9.4%, respectively, using qPCR method (Table S1). To the extent that the copy number of the 16S rRNA gene correlates with the number of cells detected by EFM quantification methods (Deng et al., 2019), the bacteria recoveries in our study were comparable to those of Highton et al. (2023). Using 16S rRNA amplicon analysis, they further showed that the bacterial community in the extracts and bulk soils shared roughly half of the ASVs and the only major difference at the family level was the greater recovery of Bacillaecae (Firmicutes) in the bulk soil. In our study, the S2/ R0 ratio for species richness of 16S rRNA and *nosZ*-I was 38.9% and 17.9%, respectively (Table S1), suggesting that roughly a quarter of the bulk soil bacteria community was extracted.

A trade-off is likely to be present between dispersion-assisted cell extraction and physical damage to bacterial cells during the dispersion especially for strongly aggregated soils. In fact, we found that greater dispersion by the sonication treatment led to the recovery of more diverse bacteria while their SAGs were, on average, lower in quality (Table 1 and Fig. S5). Genome quality is strongly affected by DNA damage caused by physical and chemical processes taking place during the sonication treatment. The sonication treatment was able to disrupt the aggregates of ca. 20-100 µm diameter sizes compared to the bead-vortexing treatment (Fig. S2). As soil bacterial community composition can be significantly different among particle/aggregate size fraction (Biesgen et al., 2020, Ranjard et al., 1998), the greater abundance and diversity of bacteria released by the sonication (Figs. 3 and 5) may be attributable to those associated with the 20-100 µm aggregates. The proportion of the total SAGs higher than the medium quality accounted for on average 16% (bead-vortexing treatment) and 5% (sonication treatment). The sonication-assisted extraction allowed the detection of *nos* and its surrounding genes even from low-quality category of SAGs (Fig. 6B). Furthermore, the sonication treatment led to higher recovery and diversity of DNA (Figs. 3 and 4A) and greater diversity of SAGs including their functional genes (Fig. 5A-C) compared to the bead-vortexing treatment. Thus, our sonication-assisted extraction method was more effective in characterizing the bacterial community present in the soil aggregates compared to the extraction with the bead-vortexing dispersion. While cavitation effect of ultrasound by sonication has been demonstrated to disrupt cell structure and associated chemical effects including the generation of free radicals can damage DNA (Li et al., 2018, Lv et al., 2019, Miller et al., 1991, Oyane et al., 2009), these negative effect may not be significant.

The current extraction method possibly led to higher recovery of soil bacterial community compared to previous soil SCG studies because no other studies used sonication to disperse soil aggregates. At the same time, the sonication-assisted extraction used in the current study likely led to greater degrees of cell damages – medium and high quality SAGs accounted for 5% (sonication treatment) and 16% (bead-vortexing treatment) of total SAGs. In comparison, Nishikawa et al. (2022) performed a mixing (details not reported) plus washing method with high-speed centrifugation for the extraction and showed that 20% of the total SAGs was in medium and high quality SAGs. Aoki et al. (2022), using a mixing pretreatment (details unknown) followed by density gradient centrifugation (Nycodenz), yielded on average 41% medium- and high-quality SAGs. These previous studies showed higher proportions of medium- and high-quality SAGs compared to the current study, which likely reflect the fact that their soil samples are typically poorly aggregated (beach, desert, mangrove, paddy soils). In our study, the dispersion using the conventional method was insufficient (Figs. S1A and S2). For a further study targeting at the bacteria in aggregates and well-aggregated soils (e.g., high in soil C, clay, and/or metal oxides), it is important to compare different methods such as the sequential washing/sonication method (Hattori, 1967, Kohno and Hashimoto, 2008) and the sequential dispersion/density gradient centrifugation method (Nadeem et al., 2013).

The physical stability of soil aggregate may exert a primary control on cell extraction efficiency given the levels of dispersive energy required for effective aggregate disruption. This hypothesis appears reasonable as cell detachment from soil surfaces is likely to be severely impeded when their habitats (e.g., pores) are present in aggregate interiors and not fully exposed to extracting solution. The aggregates studied here are relatively stable as our Acrisol is relatively high in clay and iron oxides contents (Mitsunobu et al., 2025). In these agricultural topsoils that contain significant amounts of water-stable aggregates, it would be important to sufficiently disperse aggregates for the extraction of cells that reasonably represent the soil bacterial community.

### 4.2 Greater recovery of “strongly-attached bacteria” by sonication-assisted extraction

The two aggregate dispersion techniques used in this study clearly affected the recovery of soil bacteria. The supernatant after the sonication treatment showed the bacterial community composition more similar to the residual soil microbiome compared to those after the bead-vortexing treatment (Fig. 4). The 16S rRNA amplicon sequencing results at the phylum level also showed the superiority of the sonication dispersion treatment compared to the bead-vortexing treatment but with a few exceptions. The Actinobacteriota detected after the bead-vortexing treatment was higher in relative abundance compared to the sonication treatment based on both the amplicon analysis and SAGs (Fig. 4B and Table S2). This result may be explained by the tendency that Actinobacteriota are easily dispersible (Ouyang et al., 2021) and tolerant to physical stress (Schimel et al., 2007). In contrast, the Bacteroidota were most abundant after the sonication treatment (LEfSe analysis in Table S2). Several genera of Bacteroidota are considered "hard-to-extract" taxa due presumably to their strong binding to the surfaces of soil organic and mineral particles (Ouyang et al., 2021).

The following three lines of results suggest that the sonication treatment preferentially released microbial cells that are more strongly attached to soil particle surface and/or present in the interior of aggregates compared to the bead-vortexing treatment. First, the sonication treatment liberated more *nosZ*-harboring bacteria cells and N_2_O-related functional genes compared to the bead-vortexing treatment. Specifically, a larger number of the cells containing *nosZ* -I and -II were detected after the sonication treatment (Fig. 3B, C) and the S2/ R0 ratio of *nosZ* -II (12%) was particularly higher after the sonication treatment compared to the bead treatment (1%). In addition, the sonication treatment led to a significantly higher abundance of Bacteroidota and Gemmatimonadota (Table S2), the main phylum possessing *nosZ*-II (Hallin et al., 2018). Our SCG analysis further showed that Proteobacteria and Acidobacteriota, which detected *nosZ* in this study, were more diverse with the sonication treatment (Fig. 5). Furthermore, the relative abundance of *nosZ* key denitrifying functional genes, was higher in the sonication treatment (Fig. 7).

Second, the possible localization of the extracted cells in the original aggregates can be inferred by comparing the current results with the previous study where the single aggregates (approximately 6 mm in diameter) isolated from the same bulk soil and incubated in the same way were sliced from the top to the center at ca 300 μm intervals followed by the amplicon sequencing of each slice (Mitsunobu et al., 2025). We found that two *nos*Z sequences, detected from 2 SAGs (YK46S3_sc-0263 belonging to Gemmatimonadota and YK46S3_sc-0319 belonging to Acidobacteriota), had 100% homology to the *nosZ* ASVs isolated from the aggregate interior in the previous study (Mitsunobu et al., 2025). Further analysis showed that the 2 SAGs corresponded to 2 ASVs (Gemmatimonadota) and 2 ASVs (Acidobacteriota) in the previous study (Fig. S10). These four ASVs were detected only in the interior region of the aggregates This comparison therefore supports the idea that the sonication dispersion liberated the microbial cells located in the interior of aggregate as well as its exterior.

Third, the analysis of EPS-related genes implies the preferential extraction of the cells strongly attached to soil surface and/or reside in physically-stable subunits of the aggregates. The strongly-attached bacteria are difficult to extract for two reasons. First, they are strongly attached to the surface of the soil solid phase using sticky compounds such as extracellular polymeric substances (Wagai et al., 2023), or they selectively reside in subunits of the studied macroaggregate that are difficult to break up (e.g., physically stable microaggregates present within water-stable macroaggregates, (Six et al., 2004, Yudina et al., 2022). Given that extracellular polymeric substances acts as a major binding agent for soil aggregation (Chenu and Cosentino, 2011, Costa et al., 2018, Oades and Waters, 1991), the strongly-attached bacteria are likely associated with physically-stable subunits of the incubated macroaggregates. We therefore hypothesized if the sonication treatment causes more dispersion of physically-stable aggregate subunits, the bacteria possessing EPS-related genes are released more after the sonication compared to the bead-vortexing treatment. The comparison of the functional genes related to the production of EPS (Cania et al., 2019) between the two dispersion treatments revealed that, among the 9 EPS genes detected, 3 of them were significantly higher in the sonication treatment (p < 0.05) while none was higher in the bead-vortexing treatment (Fig. S8). These differences likely resulted from non-Actinobacteriota because (i) 66 % of the bacteria extracted by SCG (Fig. 5A) and 83% of the high-quality bacteria belong to Actinobacteriota (Fig. S5), and (ii) the Actinobacteriota was characterized by a lower number of EPS genes (see Result 3.3.1) whereas these numbers were higher for the non-Actinobacteriota that were more extracted from the sonication treatment (see Result. 3.3.1.).

There are two possible explanations for the higher abundance of Actinobacteriota under the bead-vortexing treatment. First, Actinobacteriota extracted from the sonication treatment were more easily damaged than those from the bead-vortexing treatment. Second, the relative proportion of Actinobacteriota in the extracted cells was lower in the sonication treatment because the stronger aggregate dispersion by sonication released greater number of the bacteria that were strongly attached to soil particles and/or resided in more stable microaggregates, thereby diluting the Actinobacteriota population. As a further result of supporting the second, gram-positive bacteria having physical tough membrane (Schimel et al., 2007) such as Actinobacteriota, Firmicutes, and Saccharibacteria were more abundant in the bead-vortexing treatment. In contrast, gram-negative bacteria such as Proteobacteria, Acidobacteriota, Dependentiae, Verrucomicrobiota, and Bacteroidota were more abundant in the sonication treatment (Fig. 5A).

The observed differences in recovered bacteria between the two aggregate dispersion treatments may give important insights into the habitat and ecology of N_2_O reducing bacteria. Nadeem et al. (2013) distinguished two types of habitats for denitrifying microbes based on their attachment strength to soil particles using sequential dispersion/density gradient centrifugation method. For a sandy loam grassland soil, the authors showed that strongly-attached cells produced less N_2_O than loosely-attached cells and the reduced N_2_O production by the strongly-attached cells was at least partially attributable to greater N_2_O reduction (higher activity of N_2_O-reductase, *nosZ*). Thus, the strongly-attached cells were presumed to be localized at “inner” habitats such as crevices and cavities of the soil particles and/or present as persistent biofilms that are more resistant to physical dispersion (Nadeem et al., 2013). The sonication treatment in our study dispersed single aggregates more effectively into finer subunits than the bead-vortexing treatment did (Fig. S2) and released more *nosZ*-harboring bacteria (Fig. 3), implying that *nosZ*-harboring bacteria may be localized more in the aggregate subunits that are physically stable (e.g., the aggregate interiors where pore was much less abundant, Fig. S3) against the bead-vortexing and/or present in the cell attachment mode that was more susceptible to the fine air bubbles released by the sonication treatment compared to the bead-vortexing.

### 4.3 Characteristics of functional genes isolated from extracted intact cells

We were able to draw three additional inferences that were not obtainable by the metagenomics approach. For the most dominant species among the most extractable phylum, Actinobacteriota, we found inter-individual variation in the arrangement of N-cycling genes (syntenic block) within the same species. Comparison of genes around *nasC* in 4 SAGs with 100% 16S rRNA sequence match and higher (medium) genomic quality (Fig. 6A) showed that one SAG lacked the 3500 bp encoding five genes, while others did not. These results indicate that, even among very closely related SAGs, where the full 16S rRNA length is perfectly matched, genetic variations due to sequence deletions (or insertions) were present. Such deletions would be difficult to detect in MAGs constructed from shotgun metagenomic sequences as the sequences before and after the deletion are almost perfectly matched. In other words, the current study showed evidence of the variation in functional gene arrangement within one species, Actinobacteriota (i.e. even at the population level) for the first time in soil environment.

Analysis of *nosZ*-flanking regions suggested that the intact cell derived SAGs we obtained were most likely from a potentially functional *nosZ*-harboring bacteria. The genetic arrangement including *nos* cluster was similar to that of known closely related species that are shown to have *nosZ* gene expression (Fig. 6B). In particular, *Gemmatimonas aurantiaca* T-27 (Chee-Sanford et al., 2019, Park et al., 2017) has been studied in detail for N_2_O reduction. Three SAGs belonging to *Gemmatimonas* (esp. SAG of Y7B10_sc-00065) showed quite similar arrangement in the *nos* operon, indicating that the obtained SAGs are from a potentially functional *nosZ*-harboring bacteria. Therefore, not only did they retain the *nosZ*, but they were a potentially functional *nosZ*-harboring bacteria because the *nosZ* was detected as part of the *nos* operon.

The comparison of the results from the single-cell and amplicon analyses by focusing on Gemmatimonadota revealed that the sequences of *nosZ* and 16S rRNA from SCG were closer to the dominant ASV of Gemmatimonadota identified by the amplicon analyses of both *nosZ* and 16S rRNA than any of the known isolated strains in Gemmatimonadota (Fig. S9 for *nosZ* and 16S rRNA). The current study thus showed that single-cell genomics of single soil aggregate has the potential to provide genomic information on microorganisms that are closely related to those dominant in the soil aggregates based on the amplicon analysis.

### 4.4 Insights from single aggregate analyses

The rationale underlying our focus on individual soil aggregates was to achieve a more refined understanding of the taxonomic diversity and functional roles of microbial communities within intact soil microhabitats, without assuming homogeneity of bulk soil samples. We compared species and functional diversity among 6 individual aggregates based on SAG data and 11 aggregates based on amplicon data. Our results showed both uniqueness and similarity in microbial diversity and selected functional gene profiles among the studied individual aggregates.

We detected some ASVs unique to individual aggregates. Mean species accumulation plot showed the increase in observed ASVs with the increase in aggregates at least up to eighth aggregates despite that all aggregates were incubated under the same condition prior to the DNA extraction (Fig. S11A). By comparing individual aggregates and the homogenized bulk soil from the same set of soil cores, Simon et al. (2024) showed that five aggregates captured higher diversity than the bulk and the ASVs unique to these aggregates accounted for 20% of total ASVs obtained. Their results indicate that these rare species were heterogeneously distributed at the small scale (Simon et al., 2024). Similarly, we found that 27% of bacterial ASVs was unique to one of the eleven aggregates (Fig. S11B). However, on average, the unique ASVs made up for only 0.5% of the total bacterial abundances (Fig. S11B).

The comparison of N-cycling genes also showed some variations among the six aggregates. The gene families involved to anammox, nitrification, nitrogen fixation, denitrification, DNRA, were low abundant. Two of the functional genes were not detected in any aggregates (*hao*, *hzsA*). These genes showed high inter-aggregate variations. Three factors may be considered to account for the high variability of the undetected and low frequency genes. First, nitrification may be limited due to the low oxygen condition shown in our incubated aggregates (Mitsunobu et al., 2025) and the low concentration of ammonium relative to nitrate in our incubation. Second, *nifH*, *hao*, *nosZ* and *hzo* are often regarded as low abundant gene families despite their important roles in N cycling (Kuypers et al., 2018, Tu et al., 2017). Detection of low abundance genes are more difficult and subject to more errors. Third, only a small portion of the entire microbial community in each aggregate was extractable and the strongly-attached cells were likely to be less represented as discussed in 4.1 and 4.2. For example, *nosZ*-harboring bacteria, which showed increasing trend from the outer surface towards aggregate core based on qPCR (Mitsunobu et al., 2025), were undetectable in one of the three replicates in the bead-vortexing treatment (Fig. 7). In contrast, they were detected from all three replicates of the sonication treatment which led to more effective dispersion of the aggregates (Fig. S2). Furthermore, more diverse N-cycling genes (especially denitrification and DNRA) tended to be extracted after the sonication than the bead-vortexing treatment (Fig. S7), confirming the importance of the dispersion method (Discussion 4.2.). The heatmap of the relative gene abundance for each dispersion method (three boxes aligned side by side in Fig. 7 box) showed some uniqueness among the individual aggregates. The variation among the triplicates was large for *nxrAB*, *nrfA*, *nosZ* (to a limited extent, *napAB*, *nirA*, *nirKS*) in both dispersion methods, for *amoA* (to a limited extent, *norBC*) in the bead-vortexing treatment only, and for *nifH* (to a limited extetent, *narB*, *nasAB*, *nao*, *nmo*) in the sonication treatment. These inter-aggregate variation in the N-cycling genes may suggest the presence of unique microbial community structure in different aggregates (Szoboszlay and Tebbe, 2021).

The current study, nevertheless, revealed overall similarity in the bacterial community structure and their potential functions among the six aggregates, implying that single water-stable macroaggregates may represent microbially and biophysically stable units in soil. While the current study is limited in the number of cells analyzed by SCG (ca 400 per aggregate) and the extraction was conducted after the lab incubation (as opposed to freshly isolated from the soil), the three lines of evidence support this view. First, the inter-aggregate variations in the number of bacteria were relatively small both in the residues (R0) and the supernatant (S2) (quantitative PCR, Fig. 3). The variation of species richness in each fraction was also small (R0 in Table S1, S2 in Fig. 3, S3 in Fig.5B) although the effect of the two dispersion treatments was present as discussed above (section 4.2). Second, when we sum up the ASVs between the residue and supernatant (R0 and S2), more than 60% of ASVs were common across the eleven aggregates. In addition, the three most dominant phyla were also similar among the ten aggregates in R0 fraction and six aggregates in S3 fraction (Figs. 4B and 5A). Third, the bacterial functional diversity and redundancy were high in a similar degree among the six single aggregates as indicated by the comparison between the number of functional genes and SAGs (Fig. 5C) and OTUs (Fig. 5D). While such high functional redundancy of soil microbial community has been shown at bulk soil scales (Chen et al., 2021, Louca et al., 2018), our study further showed that the high redundancy is maintained even at the individual aggregate scale (Fig. 5). Fourth, we found rather similar relative abundance of the N-cycling genes across the six aggregates (Fig. 7). Of the seven major pathways we analyzed, ANR, DNRA and organic N metabolism had the highest gene abundance as depicted in the arrow thickness (Fig. 7). Our results at the aggregate level are consistent with the FACE grassland study (Tu et al., 2017) which also showed Actinobacteriota as the main phylum and organic N metabolism and nitrate reduction as the two major N transformation pathways at the bulk soil level, presumably because microbes gain energy and nutrients by these two processes (Condron et al., 2010, Moreno-Vivián et al., 1999). Furthermore, we found that the genes associated with denitrification, ANR, DNRA, and N metabolism showed high relative abundance in all six aggregates (see 3x2 box next to each arrow, Fig. 7).

Notably, all six aggregates contained microbes with genes enabling the conversion of nitrate into all possible nitrogen forms, suggesting that aggregates act as a minimal functional unit for N-cycling in soil. Similarly, Simon et al. (2024) suggested that soil aggregates may act as functional units of soil organic matter turnover based on the correlations among microbial community composition, organic matter content, and its recycling status among the intact aggregates (ca 2 mm in diameter). Similarly high functional redundancy among the aggregates shown in our study further highlights the potential value of soil aggregate as a basic experimental unit to examine microbial diversity-function relationship as aggregates largely maintain the physical, chemical, and microbial condition of *in-situ* soils. Our results pose a question regarding the efficacy of contemporary cell extraction techniques, such as bead-beating or vortexing, in soils that exhibit strong aggregation due to their suboptimal recovery of cells and DNA. It is, thus, imperative to advance extraction methodologies that will facilitate a more comprehensive understanding of the microbial diversity and functioning within soil environments.

### Data Availability Statement

The datasets, sequence raw data of SAGs and amplicon sequence data for this study can be found in the the DDBJ. All data were registered using BioProject PRJDB19546.

## Acknowledgement

The authors are grateful to S. Sato and M. Bamba (Tohoku University) for amplicon analysis, M. Hayatsu for valuable comments on extraction method development. We also thank Shizuoka Prefectural Research Institute of Agriculture and Forestry for kindly sharing soils from the long-term experimental field. This paper is based on results obtained from a project, JPNP18016, commissioned by the New Energy and Industrial Technology Development Organization (NEDO).

