## Supplement for "Single-cell genomics of single soil aggregates: methodological assessment and potential implications with a focus on nitrogen metabolism"

### Fig. S1

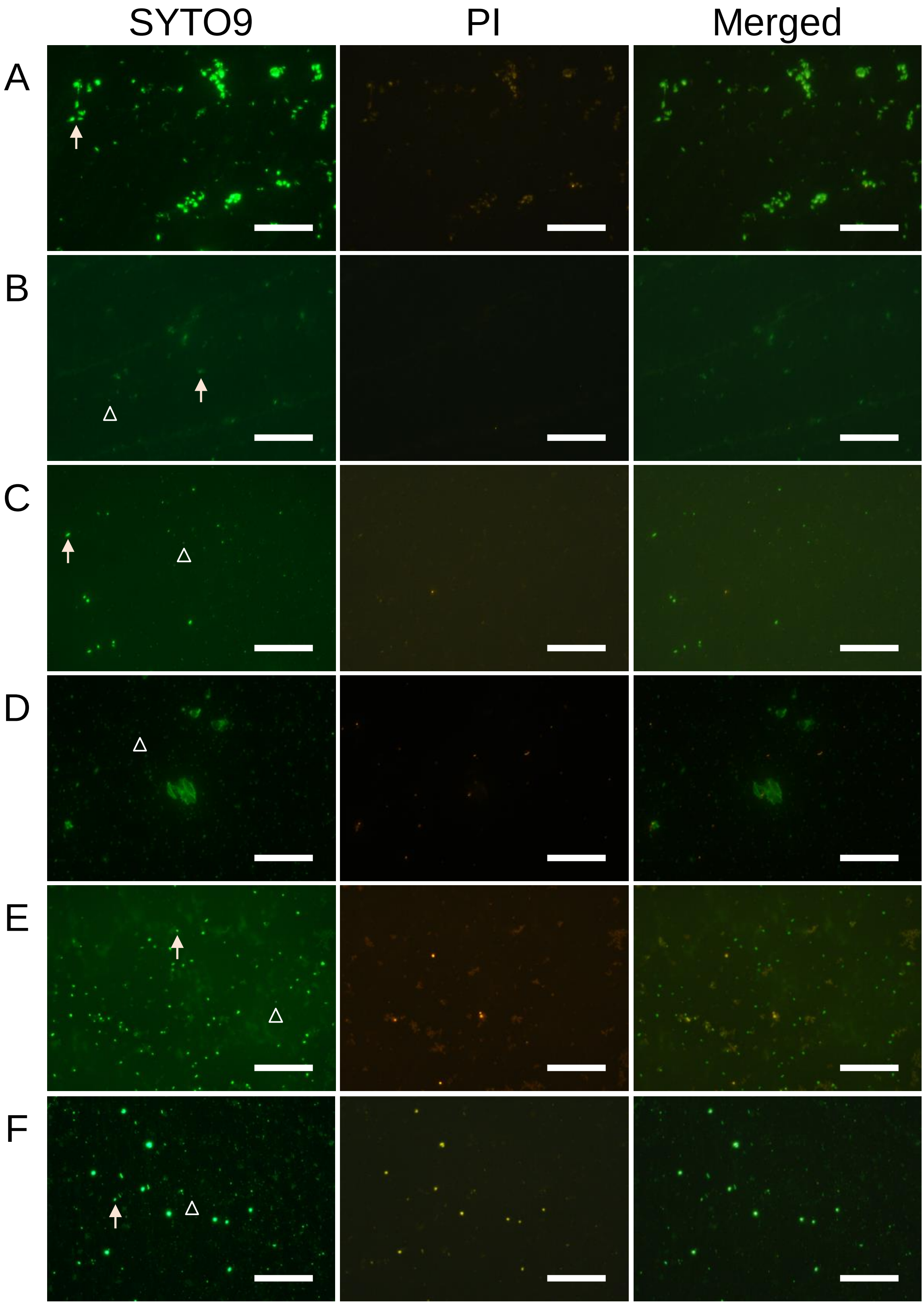


Fig. S1 Live/dead staining of the soil supernatant with conventional methods (A), with Sonic (S0, B) or Beads (S0, C), with Nycodenz purification (D), and with Sonic (S3, E) and Beads (S3, F). These samples were stained with SYTO9 and PI. SYTO9, PI and merged images were shown. Bar: 50 µm. Bacteria or soil particles were determined by the presence or absence of Brownian motion. For example, a cell similar in size to the particle indicated by the white triangle was counted as a single bacterium, whereas a cell similar in size to the particle indicated by the arrow was not counted as a single bacterium.

### Fig. S2

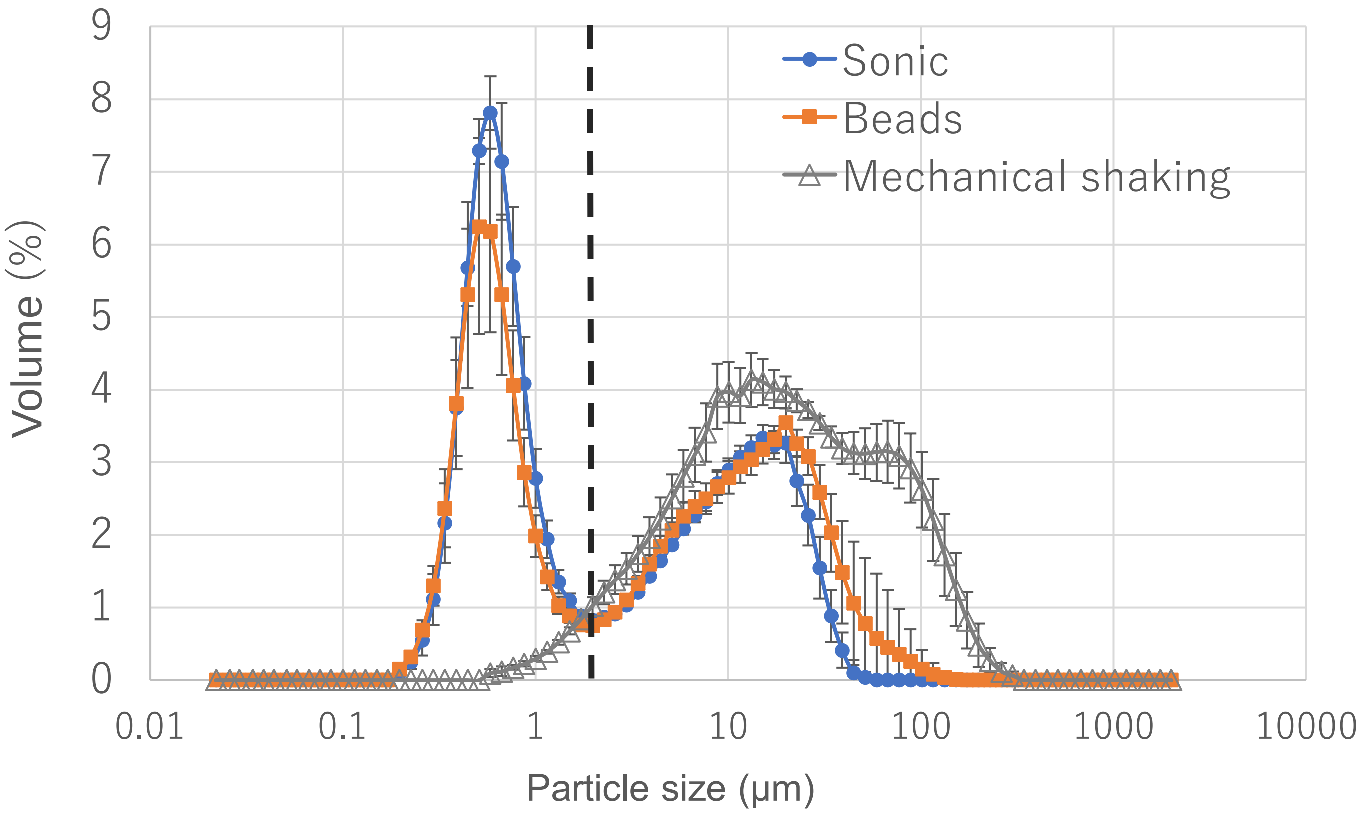


Fig. S2 Particle size distribution of single soil aggregates after the sonication, beads-vortexing and mechanical shaking dispersion treatments (n=3). The bimodal distribution boundary at 1.98 mm was shown as a vertical dotted line.

### Fig. S3

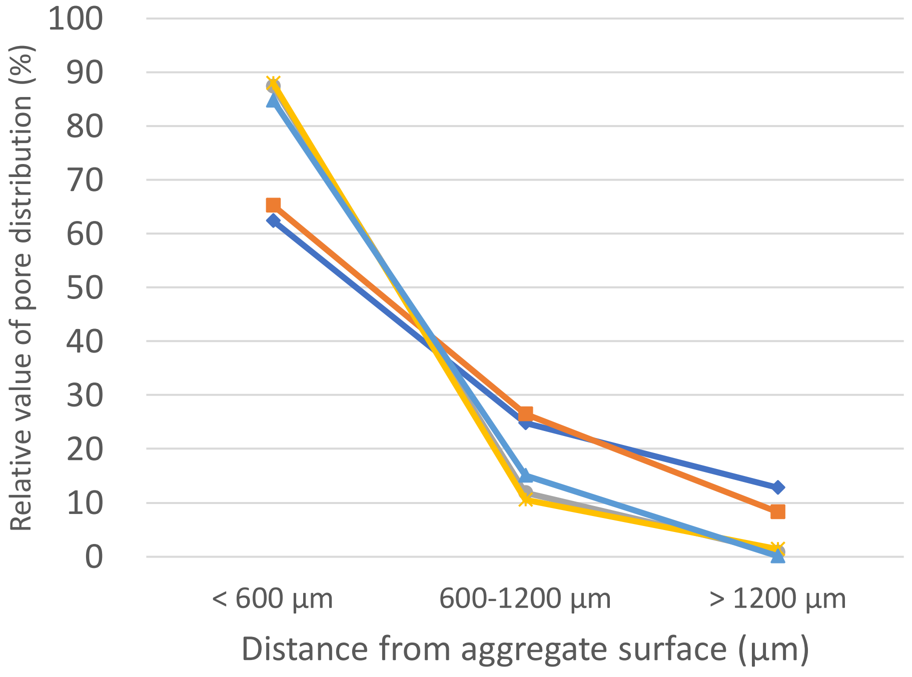


Fig. S3 Pore depth distribution of water-stable macro aggregate by X-ray µCT analysis. The different aggregates are shown by different colors and symbols.

Fig. S4

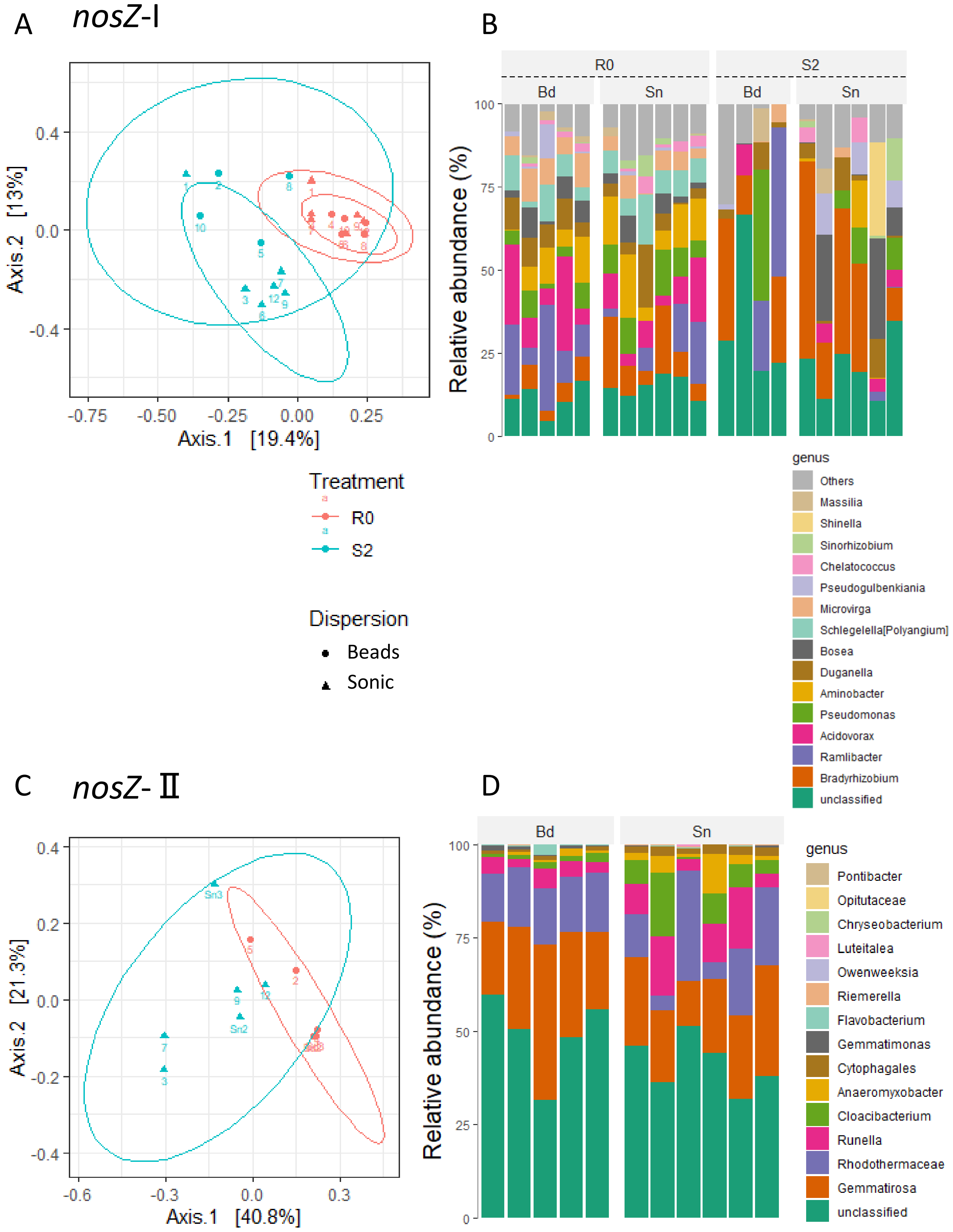


Fig. S4 Community structure by *nosZ*-I (A and B) and -II (C and D) amplicon analysis. Community differences are shown by a PCoA plot of the weighted UniFrac distance matrix (A, C) and a bar chart at the genus level (B, D) across the different fractions(R0: residue, S2: second supernatant) and dispersion methods (Bd: bead-vortexing, Sn: sonication treatment). The numbers indicate the sample ID.

### Fig. S5

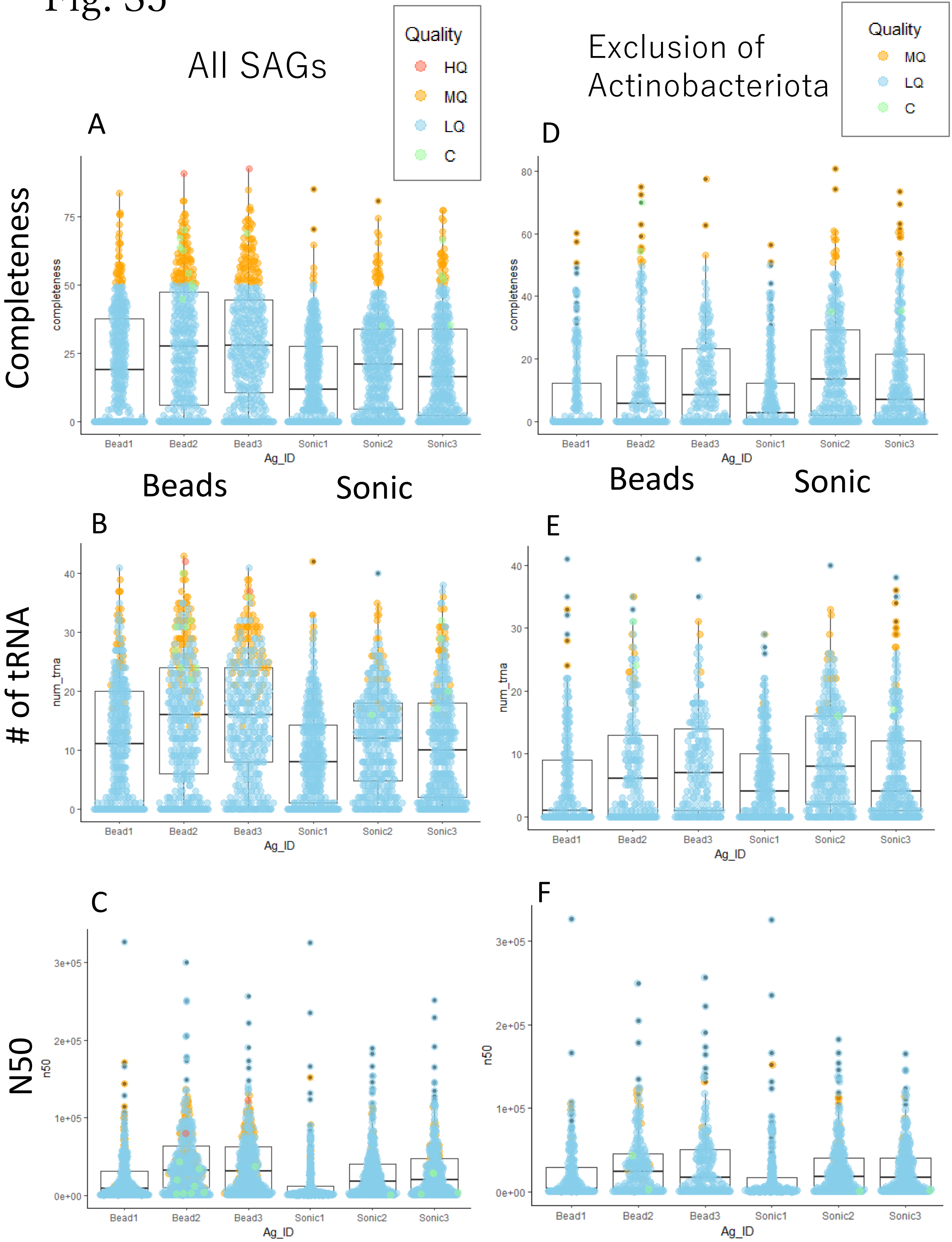


Fig. S5 Genomic quality of the SAGs. Data for all SAGs in A, B and C. Data for exclusion of Actinobacteriota in D, E and F. A and D indicates Completeness. D and E indicate number of tRNA. C and F indicate N50.

### Fig. S6

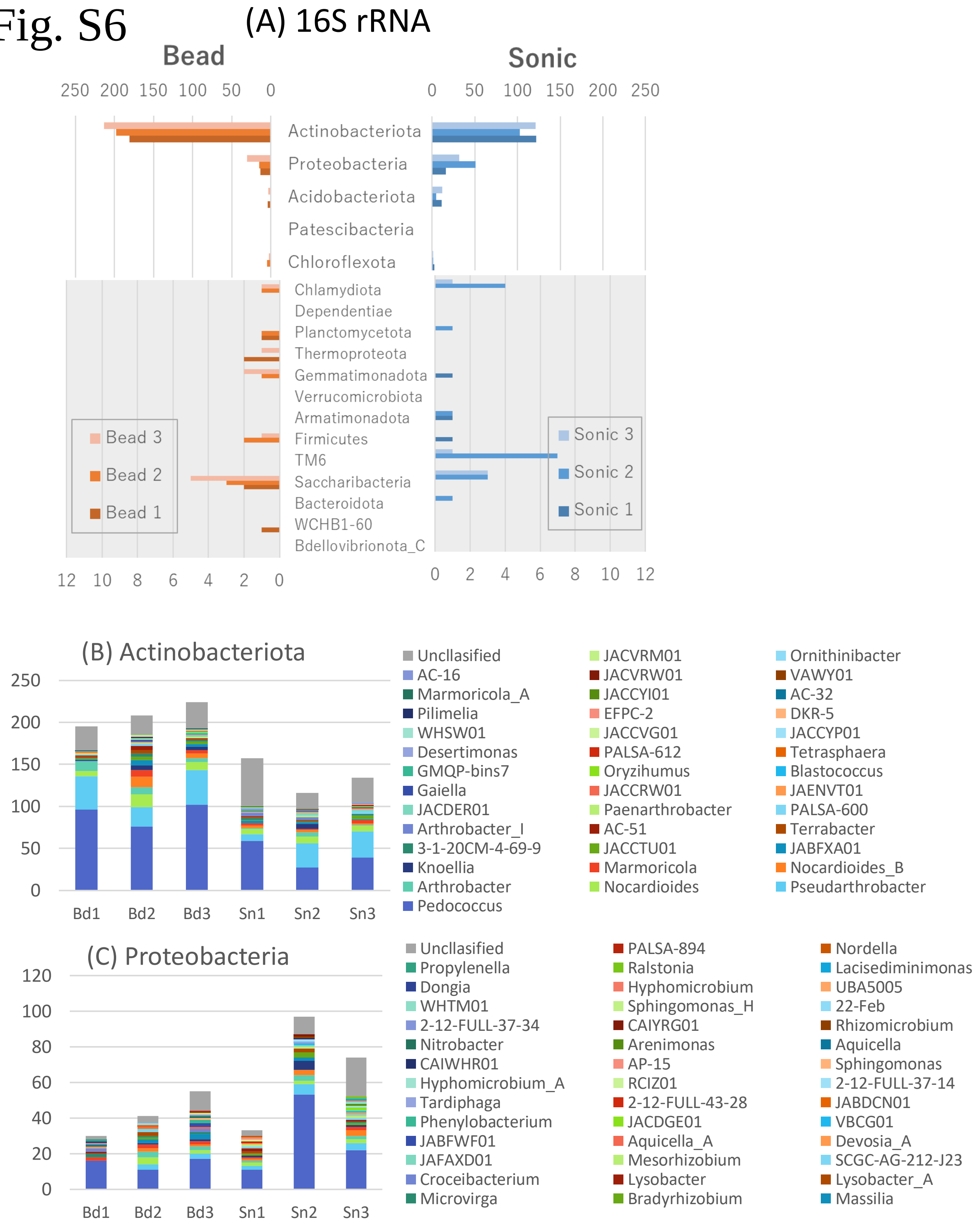


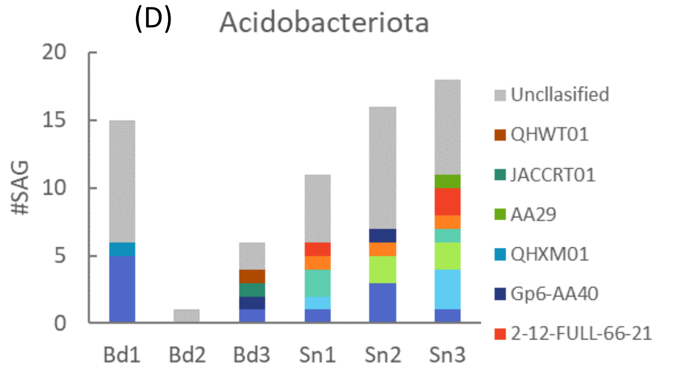


Fig. S6 Taxonomy profile of SAGs annotated by 16S rRNA region. (A) is the number of SAGs at the phylum-level for all SAGs, and (B)-(D) is the number of SAGs at the genus-level for each phylum in the top three.

### Fig. S7

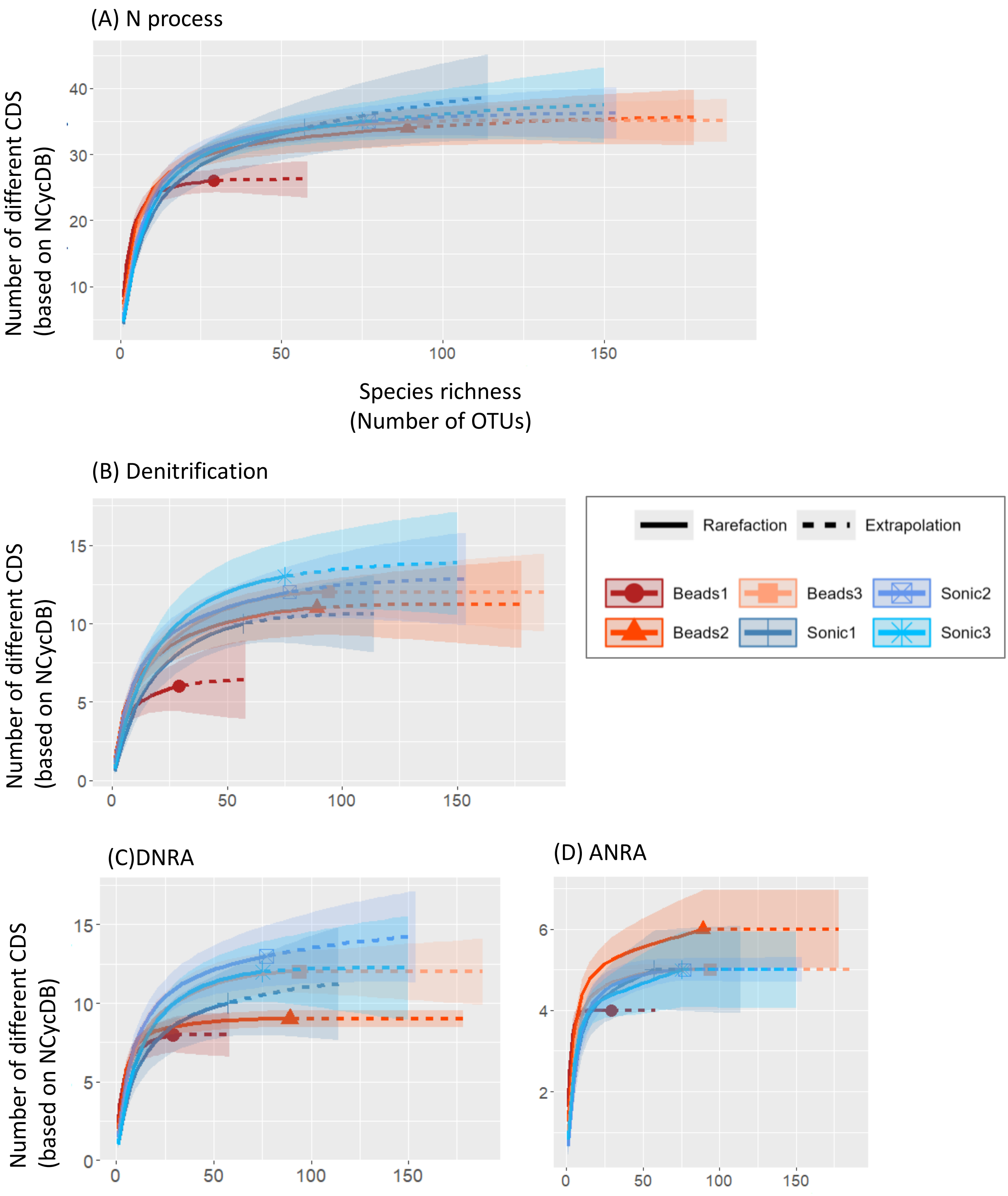


Fig. S7 Rarefaction curve of CDS related to N-cycling against number of OTUs in single aggregate.

### Fig. S8

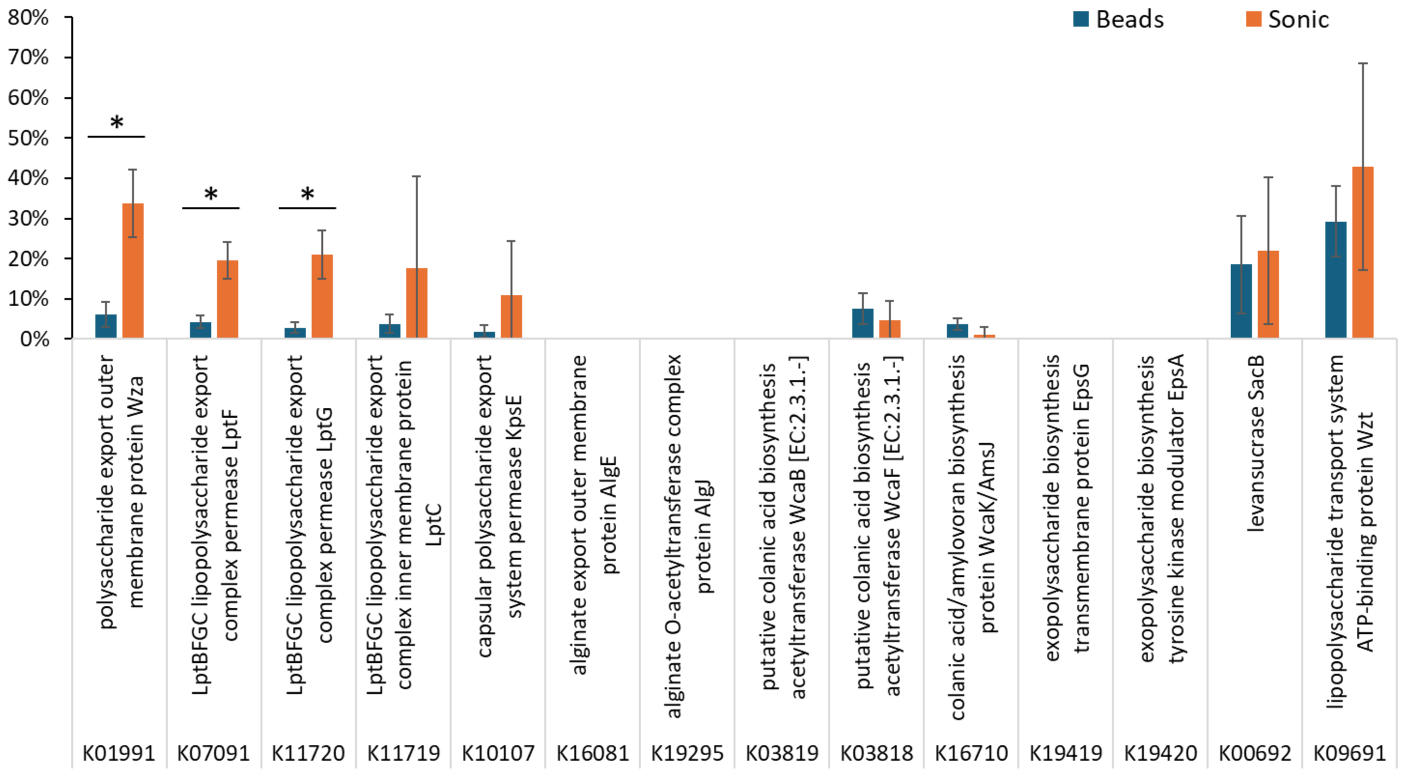


* p< 0.05, t-test, n=3, mean ± SD

Fig. S8 The proportion of SAGs harboring the EPS related genes

### Fig. S9

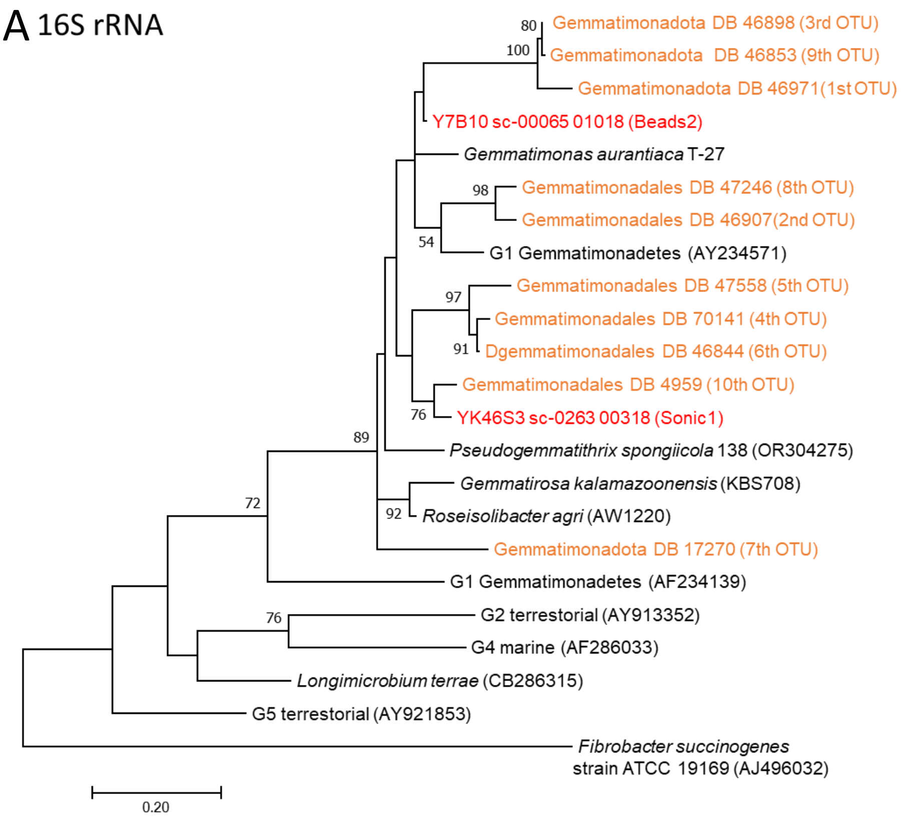


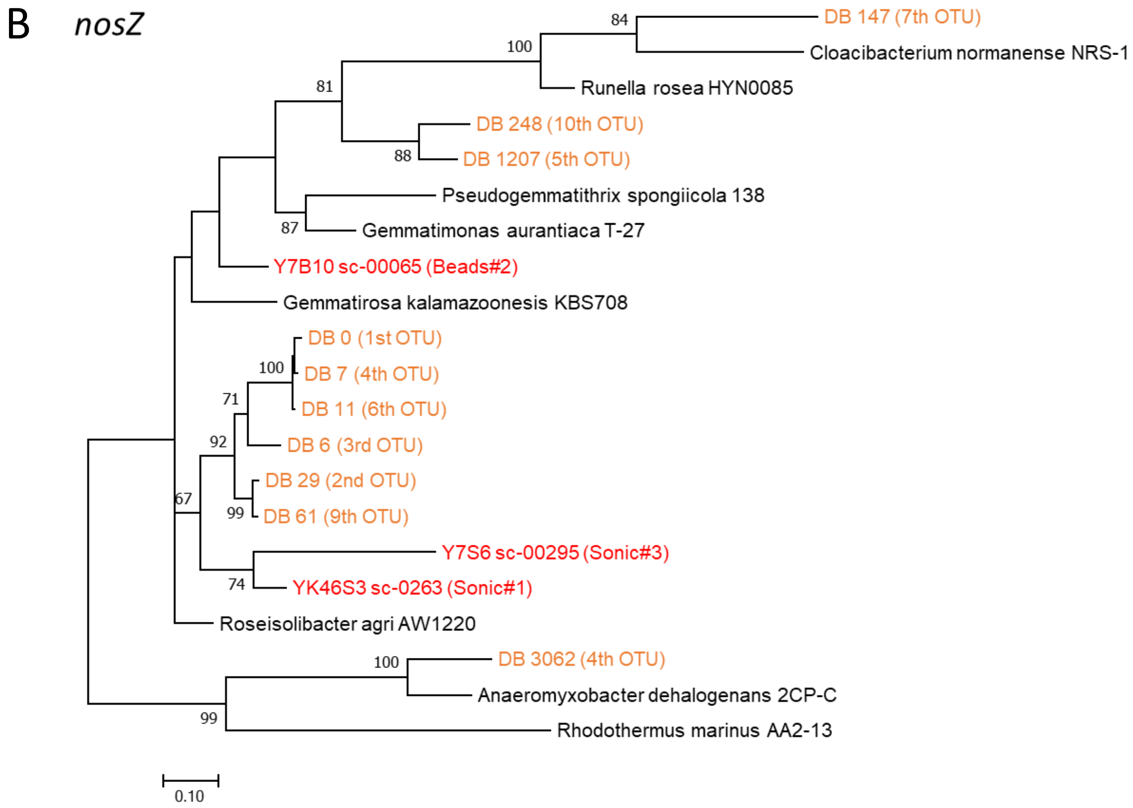


Fig. S9 Molecular phylogenetic analysis by Maximum likelihood method. A phylogenetic tree of 16S rRNA within Gemmatimonadota and *nosZ* was constructed using the top ten OTUs ( ≥99% sequence identity) in *nosZ*II amplicon analysis of the supernatant (orange color) , sequences of SAGs (red color) and reference sequences (black color). The 16S rRNA of *Fibrobacter succinogenes* strain ATCC 19169 (accession number AJ496032) as an outgroup in the phylogenetic tree (A). Bold number that support a probability >50% in bootstrap analyses (based on 500 replicates) are shown.

### Fig. S10

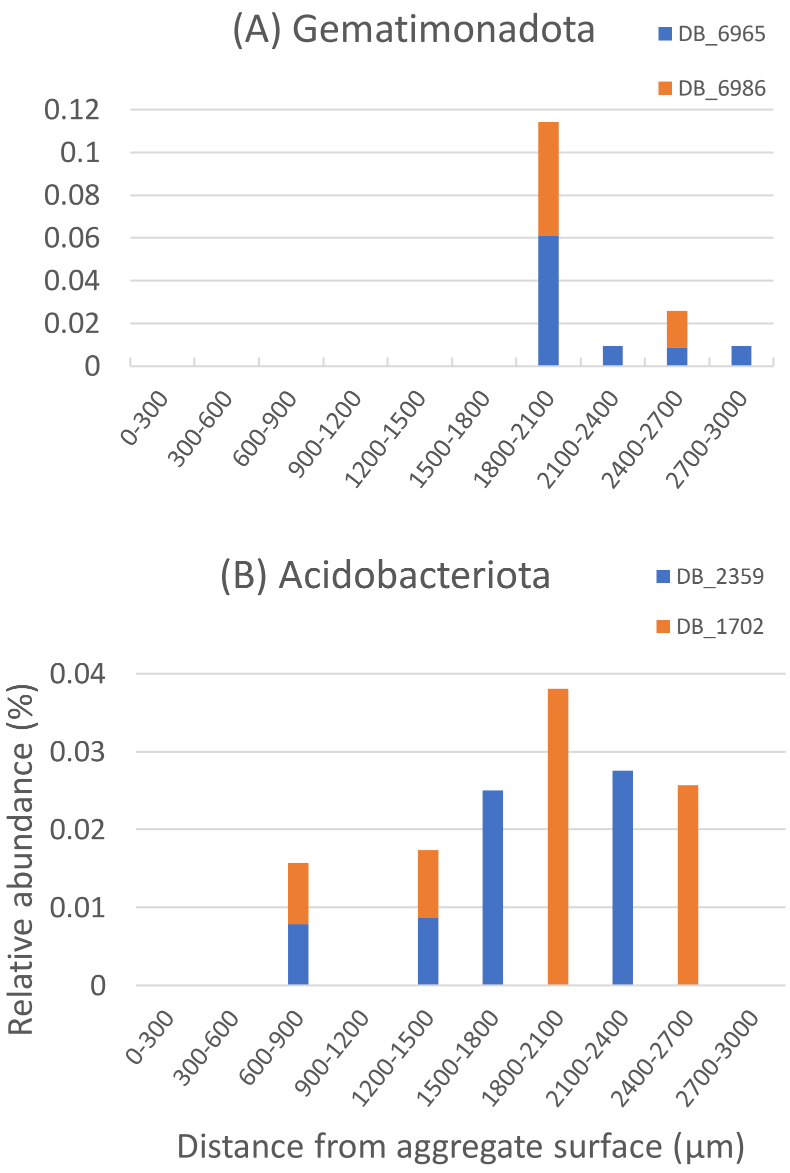


Fig. S10 The depth variations of ASVs with 100% homology to the *nosZ* of SAGs in the previous study (Mitsunobu et al. 2024).

Fig.S11

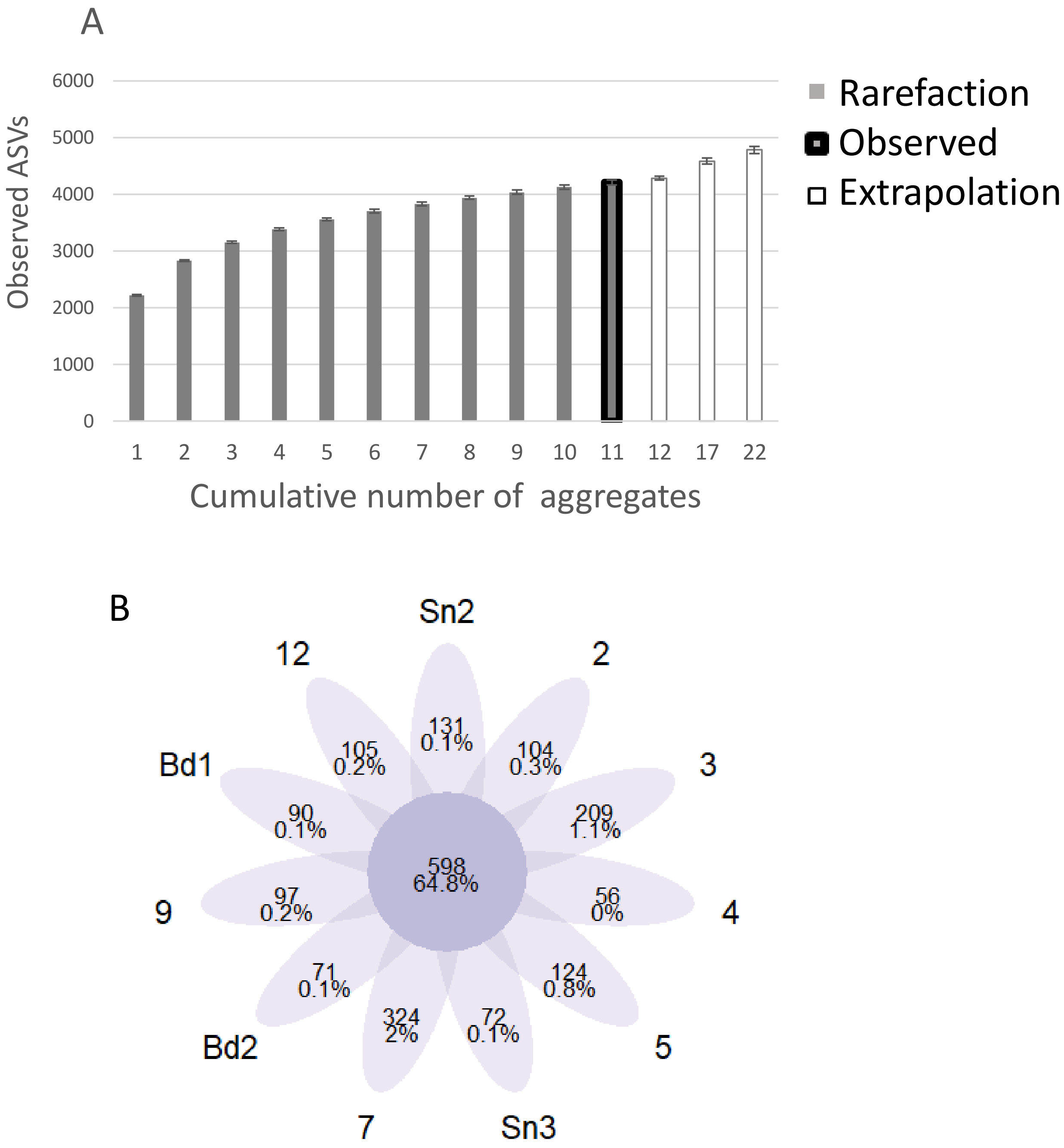


Fig.S11 Uniqueness and similarity of microbial communities of single aggregate. Species accumulation barplot (A) and venn diagram (B) were generated from ASV count tables combining R0 and S2 in each soil aggregate sample. Note that no data on S2 of aggregate #4 (missing values). The venn diagram indicated observed ASVs (the integer) and proportion of total abundances (%) of ASVs that were detected in all aggregates sample or detected exclusively only in specific single aggregate.

Table S1 Soil sample data and microbial data for each aggregate.

| | | | | | | | DNA | Copy number | | | | | |
|---|---|---|---|---|---|---|---|---|---|---|---|---|---|
| | | | | | | | | Sample | | | Per soil d. w. g | | |
| Dispersion | Fraction | Aggegate ID | Soil weight dry | Soil weight wet | Diameter | Volume | Total DNA (μg) | 16S rRNA | nosZ clade I | nosZ clade II | 16S rRNA | nosZ clade I | nosZ clade II |
| Sonic | R0 | Sn2 | 0.121 | 0.143 | 6.48 | 111.3 | 0.090 | 7.5E+06 | 1.2E+06 | 1.5E+04 | 6.2E+07 | 9.9E+06 | 1.2E+05 |
| Beads | R0 | 2 | 0.080 | 0.097 | 6.97 | 145.0 | 0.188 | 3.7E+06 | 1.2E+06 | 3.2E+04 | 4.6E+07 | 1.5E+07 | 4.0E+05 |
| Sonic | R0 | 3 | 0.054 | 0.066 | 6.13 | 98.4 | 0.074 | 1.3E+07 | 1.3E+06 | 1.0E+05 | 2.4E+08 | 2.4E+07 | 1.9E+06 |
| Beads | R0 | 4 | 0.105 | 0.128 | 6.34 | 100.0 | 0.202 | 1.0E+07 | 1.8E+06 | 1.1E+05 | 9.5E+07 | 1.7E+07 | 1.1E+06 |
| Beads | R0 | 5 | 0.055 | 0.067 | 5.78 | 77.3 | 0.043 | 4.1E+06 | 1.5E+06 | 2.9E+04 | 7.4E+07 | 2.7E+07 | 5.3E+05 |
| Sonic | R0 | Sn3 | 0.189 | 0.236 | 7.46 | 187.1 | 0.102 | 3.9E+06 | 1.5E+06 | 2.4E+04 | 2.1E+07 | 8.2E+06 | 1.2E+05 |
| Sonic | R0 | 7 | 0.052 | 0.068 | 5.71 | 71.3 | 0.100 | 4.4E+06 | 1.5E+06 | 6.8E+03 | 8.4E+07 | 2.9E+07 | 1.3E+05 |
| Beads | R0 | Bd3 | 0.168 | 0.206 | 8.21 | 190.7 | 0.388 | 4.0E+07 | 1.6E+06 | 1.2E+06 | 2.4E+08 | 9.7E+06 | 7.4E+06 |
| Sonic | R0 | 9 | 0.095 | 0.112 | 5.70 | 92.8 | 0.067 | 1.1E+07 | 1.3E+06 | 1.7E+05 | 1.1E+08 | 1.4E+07 | 1.8E+06 |
| Beads | R0 | Bd2 | 0.081 | 0.102 | 5.20 | 82.9 | 0.298 | 5.1E+07 | 1.3E+06 | 9.2E+05 | 6.3E+08 | 1.6E+07 | 1.1E+07 |
| Sonic | R0 | 12 | 0.062 | 0.073 | 6.25 | 100.6 | 0.082 | 6.6E+06 | 1.6E+06 | 1.9E+04 | 1.1E+08 | 2.5E+07 | 3.1E+05 |
| Sonic | S2 | Sn2 | | | | | | 7.8E+05 | 7.8E+04 | 3.0E+03 | 6.5E+06 | 6.5E+05 | 2.5E+04 |
| Beads | S2 | 2 | | | | | | 3.5E+05 | 3.9E+04 | 5.4E+03 | 4.4E+06 | 4.8E+05 | 6.7E+04 |
| Sonic | S2 | 3 | | | | | | 2.1E+05 | 3.4E+04 | 4.2E+03 | 3.9E+06 | 6.4E+05 | 7.9E+04 |
| Beads | S2 | 4 | | | | | | ND | ND | ND | ND | ND | ND |
| Beads | S2 | 5 | | | | | | 1.0E+05 | 3.0E+04 | 9.8E+02 | 1.8E+06 | 5.4E+05 | 1.8E+04 |
| Sonic | S2 | Sn3 | | | | | | 5.3E+05 | 1.1E+05 | 3.8E+03 | 2.8E+06 | 5.9E+05 | 2.0E+04 |
| Sonic | S2 | 7 | | | | | | 1.8E+05 | 7.7E+04 | 1.3E+03 | 3.4E+06 | 1.5E+06 | 2.4E+04 |
| Beads | S2 | Bd3 | | | | | | 2.6E+05 | 6.2E+04 | 8.1E+02 | 1.6E+06 | 3.7E+05 | 4.8E+03 |
| Sonic | S2 | 9 | | | | | | 4.6E+05 | 3.6E+04 | 2.7E+03 | 4.8E+06 | 3.7E+05 | 2.8E+04 |
| Beads | S2 | Bd2 | | | | | | 2.8E+05 | 7.3E+04 | 9.2E+02 | 3.5E+06 | 9.0E+05 | 1.1E+04 |
| Sonic | S2 | 12 | | | | | | 3.3E+05 | 2.9E+04 | 2.5E+03 | 5.3E+06 | 4.7E+05 | 4.0E+04 |
| Sonic | ND | Bd1 | 0.056 | 0.074 | 5.57 | 69.2 | ND | ND | ND | ND | ND | ND | ND |
| Beads | ND | Sn1 | 0.074 | 0.098 | 6.12 | 77.9 | ND | ND | ND | ND | ND | ND | ND |

| | | α - diversity in 16S rRNA | | | | |
|---|---|---|---|---|---|---|
| Dispersion | Fraction | Aggegate ID | Observed | Chao1 | PD | Shannon | InvSimpson |
| Sonic | R0 | Sn2 | 2241 | 2863 | 3290 | 6.15 | 123.8 |
| Beads | R0 | 2 | 1952 | 2409 | 3038 | 5.78 | 78.7 |
| Sonic | R0 | 3 | 2201 | 2865 | 3248 | 6.09 | 101.8 |
| Beads | R0 | 4 | 2218 | 2794 | 3265 | 6.16 | 135.1 |
| Beads | R0 | 5 | 1301 | 1583 | 2402 | 5.71 | 98.2 |
| Sonic | R0 | Sn3 | 1972 | 2286 | 3073 | 6.09 | 99.6 |
| Sonic | R0 | 7 | 2189 | 2650 | 3255 | 6.06 | 85.7 |
| Beads | R0 | Bd3 | 2336 | 2893 | 3383 | 6.38 | 165.0 |
| Sonic | R0 | 9 | 2195 | 2750 | 3266 | 6.24 | 147.2 |
| Beads | R0 | Bd2 | 2242 | 2803 | 3315 | 6.33 | 192.5 |
| Sonic | R0 | 12 | 2184 | 2760 | 3258 | 5.94 | 102.7 |
| Sonic | S2 | Sn2 | 1345 | 1462 | 2429 | 5.46 | 56.2 |
| Beads | S2 | 2 | 810 | 849 | 1778 | 4.89 | 27.7 |
| Sonic | S2 | 3 | 760 | 922 | 1709 | 5.96 | 194.2 |
| Beads | S2 | 4 | ND | ND | ND | ND | ND |
| Beads | S2 | 5 | 591 | 651 | 1442 | 4.83 | 21.2 |
| Sonic | S2 | Sn3 | 1117 | 1213 | 2191 | 5.44 | 60.3 |
| Sonic | S2 | 7 | 696 | 746 | 1645 | 6.18 | 314.1 |
| Beads | S2 | Bd3 | 881 | 918 | 1890 | 4.47 | 19.1 |
| Sonic | S2 | 9 | 1090 | 1161 | 2159 | 5.47 | 49.4 |
| Beads | S2 | Bd2 | 866 | 920 | 1872 | 4.59 | 17.2 |
| Sonic | S2 | 12 | 1074 | 1160 | 2128 | 5.54 | 69.4 |
| Sonic | ND | Bd1 | ND | ND | ND | ND | ND |
| Beads | ND | Sn1 | ND | ND | ND | ND | ND |

Table S2 The biomarker that characterizes the dissimilarity of the community at the phylum level among treatments by LEfSe analysis."Group" includes the residue after the beads treatment (R0Bd) and the sonication treatment (R0Sn), the supernatant after the beads treatment (S2Bd) and the sonication treatment (S2Sn).

| Domain | Phylum | Group | LDA | P.unadj | P.adj | Significance |
|---|---|---|---|---|---|---|
| Bacteria | Chloroflexota | R0Bd | 3.625 | 0.008 | 0.034* | |
| Bacteria | Proteobacteria | R0Bd | 5.243 | 0.001 | 0.014* | |
| Bacteria | Nitrospirota | R0Bd | 3.569 | 0.019 | 0.056 | |
| Archaea | Euryarchaeota | R0Sn | 2.323 | 0.002 | 0.016* | |
| Bacteria | Fibrobacterota | R0Sn | 2.974 | 0.000 | 0.014* | |
| Bacteria | Firmicutes | R0Sn | 2.859 | 0.001 | 0.014* | |
| Bacteria | Actinobacteriota | S2Bd | 5.289 | 0.010 | 0.037* | |
| Bacteria | Parcubacteria | S2Bd | 3.390 | 0.011 | 0.040* | |
| Bacteria | Armatimonadota | S2Sn | 3.338 | 0.005 | 0.027* | |
| Bacteria | Bacteroidota | S2Sn | 3.315 | 0.004 | 0.022* | |
| Bacteria | Candidatus_Saccharibacteria | S2Sn | 4.409 | 0.001 | 0.014* | |
| Bacteria | Gemmatimonadota | S2Sn | 3.876 | 0.003 | 0.018* | |
| Bacteria | Chlamydiota | S2Sn | 3.615 | 0.026 | 0.070 | |
| Bacteria | Candidatus Latescibacterota | S2Sn | 2.823 | 0.037 | 0.090 | |
